# A facile chemical strategy to synthesize precise AAV-protein conjugates for targeted gene delivery

**DOI:** 10.1101/2024.07.20.604406

**Authors:** Quan Pham, Jake Glicksman, Sadegh Shahraeini, Boyang Han, Delilah Jewel, Conor Loynd, Souyma Jyoti Singha Roy, Abhishek Chatterjee

## Abstract

The efficacy of current gene therapy approaches using adeno associated virus (AAV) vectors is limited by the poor control over their tissue tropism. Untargeted AAV vectors require high doses to achieve therapeutic efficacy, which is associated with toxic off-target impacts and increased therapeutic costs. The ability to reprogram existing AAV vectors to selectively transduce target tissues is essential to develop next-generation human gene therapies that are safer, more efficacious, and less expensive. Using selective and high-affinity antibodies and antibody-like proteins to retarget existing AAV vectors to bind novel cell-surface receptors offers an attractive and modular approach to reprogram their tropism. However, attaching these proteins onto the complex and delicate AAV capsids remains challenging. Here, we report a versatile chemical strategy to covalently attach recombinant proteins onto the capsid of AAV, using a combination of genetic code expansion and bioorthogonal conjugation chemistry. This method is efficient, and allows precise control over the site and stoichiometry of protein attachment onto the AAV capsid, enabling systematic optimization of the resulting conjugate. Using this approach, we generated conjugates of AAV2 with an anti-HER2 nanobody and a full-length anti-HER2 IgG, which show highly efficient and selective gene delivery into HER2+ cancer cells. Remarkably, the optimized AAV2-nanobody conjugate facilitated efficient transduction of HER2+ tumor xenograft in mice with little off-target gene expression, including in the liver. Programmable synthesis of AAV-protein conjugates using this method offers a promising new strategy to rationally engineer next-generation gene therapy vectors.

## Introduction

Adeno-associated virus (AAV) mediated delivery of therapeutic genetic cargo *in vivo* has advanced at a rapid pace in recent years, enabling first-in-class human gene therapies such as Luxturna, Zolgensma, and Hemgenix.^1–5^ The remarkable success of AAV vectors can be attributed to their unique features such as non-pathogenic nature, low immunogenicity, and the ability to provide long-term gene expression.^1–3^ However, our lack of control over the tropism of available AAV vectors remains a key challenge for developing next-generation gene therapies with superior efficacy and safety profiles. Due to the current reliance on native AAV serotypes, which exhibit suboptimal tissue specificity, very high doses are typically required to achieve sufficient on-target delivery, which leads to unintended off-target effects and high costs of treatment.^1–6^

Numerous strategies have been explored to improve the tissue-specificity of AAV vectors,^4^ which broadly falls into two categories: directed evolution based approaches, and rational incorporation of known targeting elements onto the virus capsid. Directed evolution based methods typically rely on generating a large library of capsid variants through different strategies, including error-prone mutagenesis,^7–9^ DNA-shuffling of existing capsid variants,^10–13^ or the insertion of a randomized peptide loop at permissive sites within the capsid protein,^14–21^ followed by their selection *in vitro* or in live animals for desired tropism. This approach has been used to develop several AAV mutants that exhibit improved selectivity for transducing specific types of target cells. Nonetheless, directed evolution of AAV capsids is a time-consuming and resource-intensive process, and an optimal outcome is not always guaranteed. Additionally, the improved properties of the resulting capsid variants do not always translate across preclinical models.

Attaching a known targeting element onto the virus capsid, which can selectively bind a cell-surface receptor expressed on the target tissue, has also been used as an alternative method for retargeting existing AAV vectors. Small retargeting ligands, such as short peptides and aptamers have been used for such proof-of-concept applications,^4, 22–25^ but the clinical utility of these approaches are limited by suboptimal affinity, selectivity, and *in vivo* stability of such ligands. Antibodies and similar proteins (e.g., affibodies) can be readily evolved to bind desired cell-surface receptors with excellent affinity and selectivity.^26–28^ These proteins have been used with much success in clinical settings for targeted delivery of other classes of therapeutics, such as antibody-drug conjugates and bispecific antibodies.^29, 30^ These proteins are particularly attractive candidates for retargeting AAV, but displaying them on the complex and delicate virus capsid is non-trivial due to perturbations of capsid assembly and packaging. It has been possible to fuse small antibody-like proteins onto the N-terminus, or at an internal loop of minor capsid proteins of AAV.^31–37^ However, only small and well-folded proteins are compatible with such fusion, and even then, it can disrupt capsid architecture and packaging efficiency. It has also been possible to attach proteins onto the AAV capsid using HUH tag and DNA linkers.^38^ However, the use of the bacteria-derived HUH protein and DNA as a linker raise major concerns over the immunogenicity and stability of such conjugates *in vivo*.

Post-packaging chemical labeling of the virus capsid is an attractive alternative for attaching retargeting groups, since it does not disrupt the virus assembly process.^4^ Native amino acids such as lysine, arginine, and tyrosine,^25, 39–41^ as well as noncanonical amino acids with bioorthogonal conjugation handles^22–24, 42–47^ have been used to functionalize the AAV capsids for various applications, including retargeting. However, linking together the AAV capsid and a protein through chemical conjugation remains challenging due to the slow reaction kinetics of traditional bioorthogonal conjugation reactions.^48, 49^ Here we report a general strategy to efficiently generate any AAV-protein conjugate with unparalleled control over the attachment sites and the stoichiometry of capsid labeling. Using the genetic code expansion (GCE) technology,^50–52^ a noncanonical amino acid (ncAA) with a bioorthogonal conjugations handle can be site-specifically introduced onto the AAV capsid as well as in the recombinant protein with complete site control. Next, the recombinant protein can be covalently attached onto the AAV capsid using a bifunctional linker using an ultra-fast bioconjugation chemistry. Using this strategy, we generated AAV2 conjugates with anti-HER2 nanobody as well as a full-length anti-HER2 antibody, which exhibit highly efficient and selective transduction of HER2+ cell lines. The AAV2-nanobody conjugate was systematically optimized by exploring various sites of attachment, linker length, and stoichiometry of labeling. The optimized conjugate enabled transduction of HER2+ tumor xenografts in mice with remarkable efficiency and specificity, showing little off-target infectivity, including in the liver. The ability to attach any recombinant protein onto the AAV capsid using this method will unlock numerous powerful opportunities to engineer its properties to develop next-generation gene-therapy vectors.

## Results and discussion

### Efficient conjugation of recombinant proteins with precise site-control

We started by exploring chemistries that will be suitable for efficient generation of AAV-protein conjugates. Linking together large proteins using chemical conjugation has been traditionally challenging due to the intrinsic deceleration of reaction kinetics for these macromolecules, as well as the practical limits on the upper limit of their concentrations that can be used.^48^ As a simple model system for screening various conjugation chemistries, we chose the crosslinking of recombinant sfGFP-151-AzK, which displays the azide-containing noncanonical amino acid AzK at the surface exposed 151 position (Figure 1a) for facile bioorthogonal labeling. The AzK residue was functionalized with different bifunctional linkers for subsequent protein-protein crosslinking. First, we probed the efficiency of crosslinking sfGFP-151-AzK using a DBCO-PEG4-DBCO linker (Figure 1b). The strain promoted azide-alkyne cycloaddition (SPAAC) has been used in the past to link proteins together.^53^ Excess DBCO-PEG4-DBCO was first used to label sfGFP-151-AzK to generate **2** (now displaying a DBCO functionality), which was purified by removing the excess linker, and incubated with fresh sfGFP-151-AzK in equimolar ratio to generate the target protein-protein conjugate **3** (Figure 1b). Even though the formation of the expected product was observed by SDS-PAGE (Figure 1c), it was slow and remained far from complete even after prolonged incubation. We concluded that the reaction kinetics of SPAAC is not well-suited for efficiently generating protein-protein conjugates.

**Figure 1:**
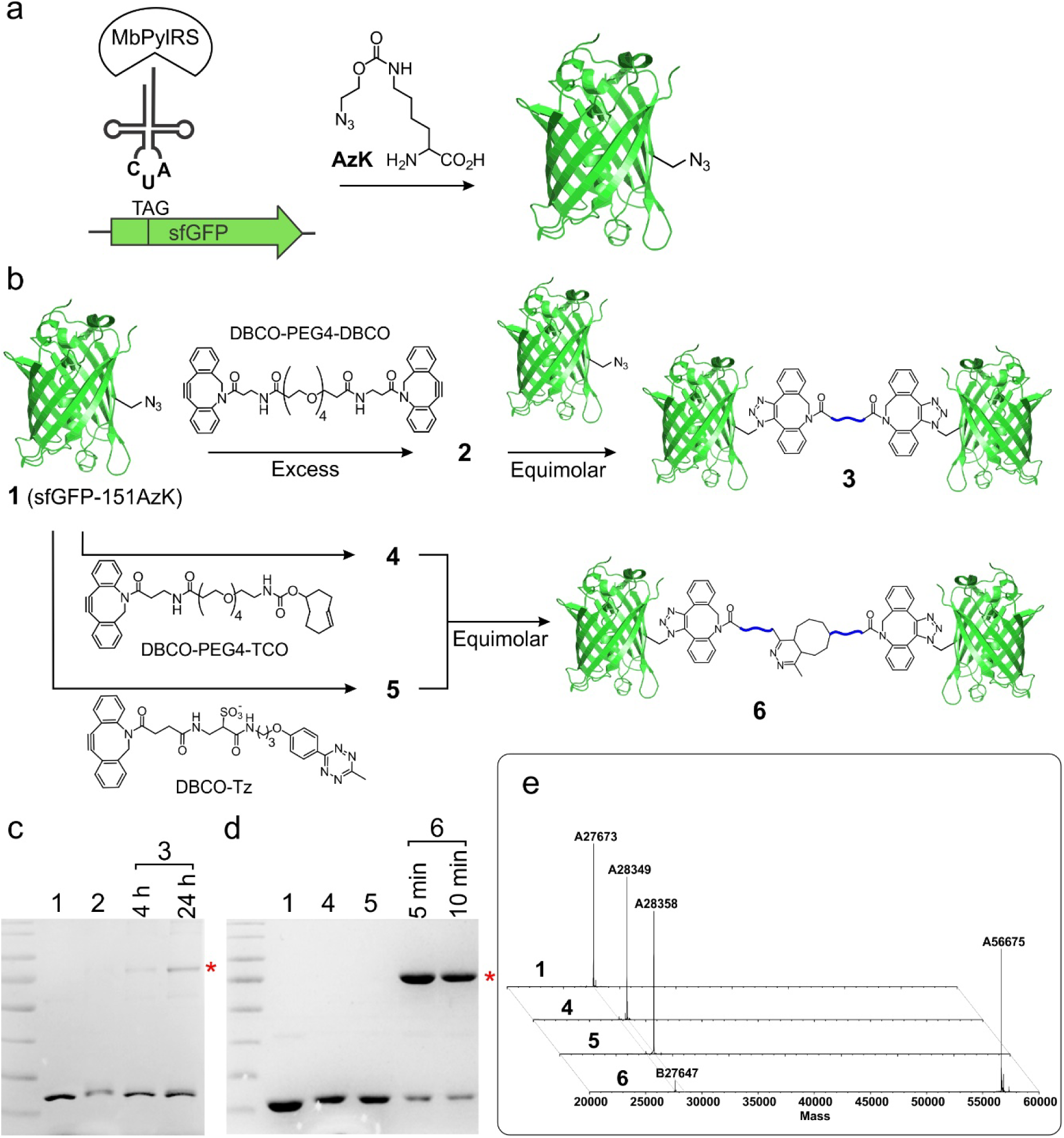
Site-specific protein-protein conjugation using GCE and bioorthogonal chemistry. a) Strategy for site-specific incorporation of AzK into recombinant proteins such as sfGFP using the PylRS/tRNA pair. b) Approaches to conjugate two sfGFP-151-AzK molecules through SPAAC or IEDDA using bifunctional linkers. c) SDS-PAGE analysis of the SPAAC-mediated conjugation reaction to generate **3**; the asterisk highlights the putative coupling product. d) SDS-PAGE analysis of the IEDDA-mediated conjugation reaction to generate **6**; the asterisk highlights the putative coupling product. e) Deconvoluted MS analysis of sfGFP-151AzK **1**, the intermediate derivatives **4** and **5**, and the final coupling product **6**, as described in panel b.

Inverse electron demand Diels-Alder (IEDDA) reaction has recently emerged as a faster alternative for bioorthogonal conjugation.^54^ We reasoned that it may be better suited for the efficient synthesis of protein-protein conjugates. This strategy has indeed been used to generate protein-protein conjugates in recent years.^48, 54–56^ To explore this possibility, we functionalized sfGFP-151-AzK separately with two synthetic bifunctional linkers, DBCO-Tz (tetrazine) and DBCO-PEG4-TCO (trans-cyclooctene), to generate conjugates **4** and **5**, site-specifically displaying the Tz and TCO functionalities, respectively (Figure 1b). After removing the excess linker, we incubated **4** and **5** in equimolar ratio, which resulted in a remarkably rapid formation of the cross-linked product **6** within minutes in excellent yields (>80%). The formation of **6** was confirmed by both SDS-PAGE analysis, which revealed the appearance of the cross-linked product as a higher molecular-weight band (Figure 1d), as well as by whole-protein ESI-MS analysis (Figure 1e, S1). The reaction was found to be nearly complete; a small amount of remaining monomeric sfGFP (<20%) was due to the reduction of the azide functionality (Figure 1e, S1).

It has been possible to genetically encode ncAAs that contain tetrazine and TCO functionalities,^57–59^ which provide the opportunity to directly introduce these groups into recombinant proteins in a site-specific manner. However, our two-step approach, involving co-translational incorporation of AzK followed by the post-translational attachment of TCO or tetrazine through SPAAC, offers a few advantages: A) The incorporation efficiency of AzK in mammalian cells is significantly higher relative to ncAAs with TCO and tetrazine groups, which will minimize any yield loss with ncAA incorporation;^60^ B) AzK is well-tolerated at numerous positions on the AAV capsid,^4, 22, 23^ whereas incorporation of other bulkier ncAAs in this context has been considerably more challenging; C) Unlike directly linking the AAV capsid and the recombinant, the use of a bifunctional linker provides additional opportunity to fine-tune the properties of the conjugate by manipulating the length and the chemical features of the linker.

### A facile strategy to generate AAV-protein conjugates

Next, we tested the feasibility of using this strategy to link together AAV and recombinant proteins. Previously, we and others have used the pyrrolysyl-tRNA synthetase (PylRS)/tRNA pair to site-specifically incorporate AzK into various positions of the AAV capsid,^4, 22–24, 42–47^ and functionalized the resulting azide handle for capsid labeling by SPAAC. Furthermore, we recently developed technology for selectively incorporating AzK into any subset of AAV capsid proteins – such as VP1-only or VP2-only (5 copies per capsid each) or VP1+VP2 (10 copies per capsid) – gaining control over the number of AzK residues introduced per capsid.^23^ The ability to systematically optimize the site and stoichiometry of capsid modification in this manner helped us demonstrate that the retargeting of AAV using a small peptide ligand worked best when a limited number of peptide ligands (approximately 10) are introduced per capsid; higher degrees of capsid modification were found to be detrimental, progressively diminishing infectivity. Guided by these insights, we chose to focus on attaching recombinant proteins only onto the minor capsid protein VP1 of AAV (5 per capsid) to avoid perturbations from excessive labeling.

AzK was selectively incorporated into the minor capsid protein VP1 using a 3-plasmid transient transfection system we previously developed (Figure 2a).^22, 23^ Briefly, expression of VP1 was uncoupled from the AAV2 *Cap* gene by mutating the start codon from ATG to CTC; it was expressed *in trans* driven by a CMV promoter. The surface-exposed site 454 in VP1 was mutated to UAG, which was suppressed using a mammalian-cell optimized MbPylRS/tRNA pair with AzK.^61, 62^ Production of AAV2 incorporating AzK at site 454 of VP1 (termed VP1-454AzK) was performed as described previously.^23^ The packaging yields (genomic titer) and infectivity of VP1-454AzK were comparable to the wild-type (WT) virus (Figure 2b, 2c, S4). During the packaging of VP1-454AzK, when AzK was omitted from the medium as a negative control, we still observed robust virus packaging (by titering packaged genomes), but the resulting virus was non-infective (Figure 2c, S4). This is because capsids with only VP2 and VP3 can still form in the absence of VP1 expression, but the resulting capsids are not infective due to the essentiality of VP1 during cellular entry for endosomal escape.

**Figure 2:**
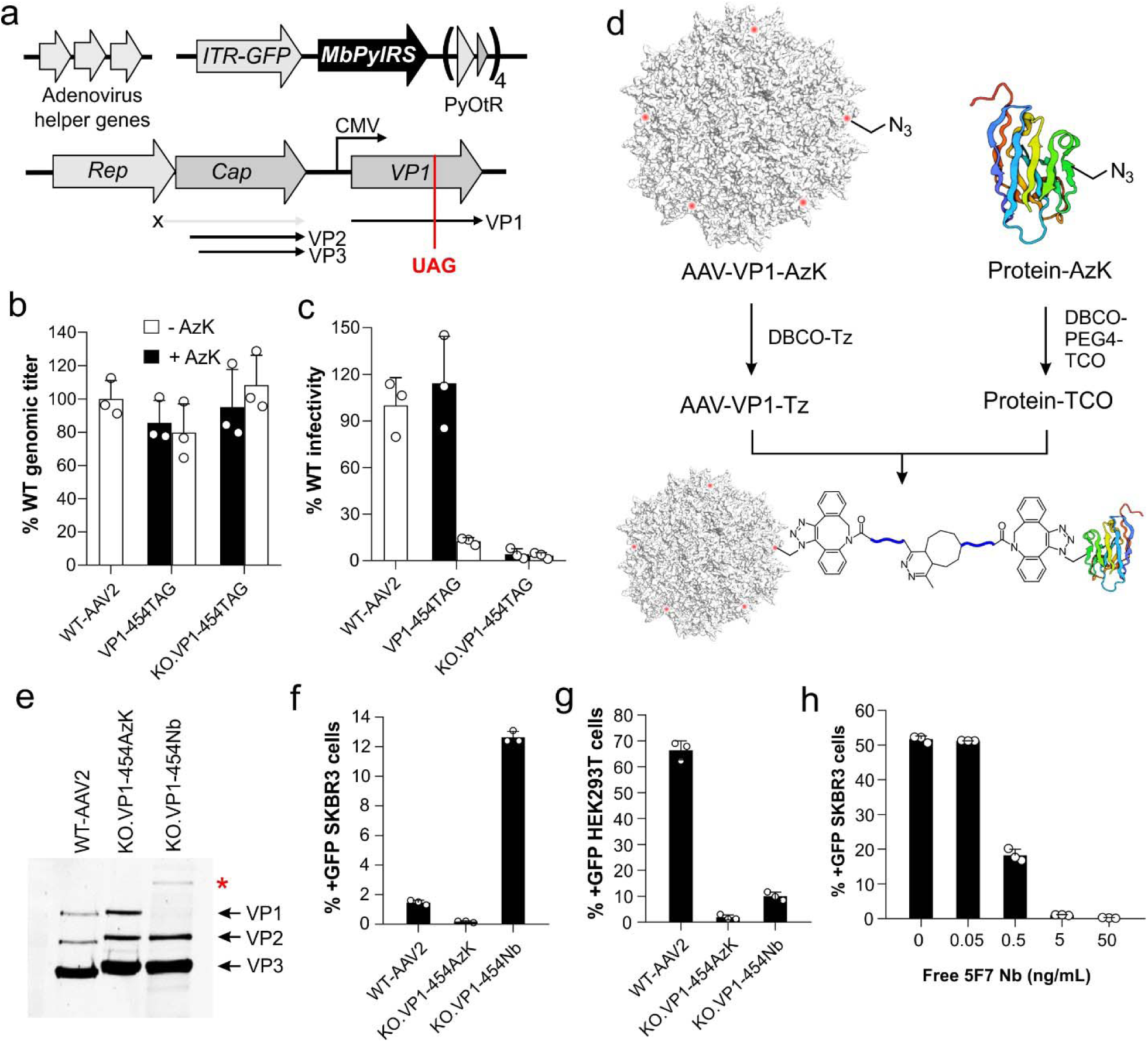
Generation of AAV-protein conjugates. a) The scheme for producing AAV2 site-specifically incorporating AzK within the VP1 minor capsid protein (∼5 copies per capsid), using a 3-plasmid transfection system. Genomic titer (b) and infectivity (c) of AAV2 (WT and KO), containing AzK at site 454 in VP1, in the presence or absence of 1 mM AzK in the media. Packaged genome titer was measured by qPCR and was normalized to WT AAV2 titer. Infectivity was measured using the characteristic fluorescence from the expression of an encoded EGFP reporter, upon infecting HEK293T cells at a constant MOI of 50, and was normalized to WT AAV2. d) The strategy to conjugate AAV2 and a recombinant protein, each containing site-specifically incorporated AzK residues. e) SDS-PAGE analysis of the conjugation reaction between DBCO-Tz-functionalized KO.VP1-454AzK, and DBCO-PEG4-TCO-functionalized Nb-69-AzK conjugation. The asterisk highlights the band corresponding to the VP1-Nb conjugate. f) Infectivity of WT, KO.VP1-454AzK, and KO.VP1-454Nb, measured by the percentage of positive GFP cells by flow cytometry, upon infecting SK-BR-3 (HER2+) cells at a constant MOI 125. g) Infectivity of WT, KO.VP1-454AzK, and KO.VP1-454Nb, measured by the percentage of positive GFP cells by flow cytometry, upon infecting HEK293T cells (low HER2 expression) at a constant MOI of 50. h) Free 5F7 nanobody inhibits the infectivity of KO.VP1-454Nb toward SK-BR-3 cells in a dose-dependent manner. Infectivity was measured by infecting SK-BR-3 cells with a constant MOI (500) of KO.VP1-454Nb, in the presence of increasing concentration of free nanobody. All data shown as the mean ± s.d. of *n* = 3 replicates.

To link together sfGFP to AAV through IEDDA as a model system, we first functionalized VP1-454AzK and sfGFP-151-AzK with DBCO-Tz and DBCO-PEG4-TCO, respectively (Figure S5a). Incubation of Tz-functionalized AAV with 1.0 μM of TCO-functionalized sfGFP for 4 h resulted in efficient formation of the AAV-sfGFP conjugate through the VP1 capsid protein, as shown by a near-complete upward shift of the VP1 band in SDS-PAGE analysis. The workflow described here provides a straightforward method to link recombinant proteins (Figure S5b) onto AAV capsids.

### Site-specific AAV2-nanobody conjugates retargeted to the HER2 receptor

The ability to seamlessly conjugate recombinant proteins onto the AAV capsid using the method described above offers the opportunity to rationally engineer its tropism through the attachment of a retargeting antibody fragment. To explore this possibility, we selected the HER2 receptor, which is overexpressed on a variety of cancer cells, and which can be selectively and efficiently targeted using well-established antibodies, nanobodies, etc.^63^ As a proof-of-concept, we selected the previously reported HER2-targeting nanobody (Nb; clone 5F7) for attachment onto the AAV capsid.^64^ 5F7 was mutated at the permissive 69 site to introduce a UAG codon, and the resulting construct was expressed in *E. coli* to incorporate AzK using an orthogonal MmPylRS/tRNA pair. Nb-69-AzK was expressed in excellent yields (>50% of WT) and MS analysis confirmed successful incorporation of AzK at the intended site (Figure S2).

Before retargeting AAV2 to a new receptor, we first eliminated its natural tropism by ‘detargeting’ it from its native primary receptor, heparan sulfate proteoglycan (HSPG).^22, 23^ To this end, we mutated two key HSPG-binding residues (R585A and R588A) on all capsid proteins, yielding a non-infective capsid termed KO.^22, 23^ AzK was selectively incorporated at site 454 of VP1 within the KO capsid, using the same strategy described above for WT-AAV2 (Figure 2b, 2c, S4), to generate KO.VP1-454AzK.

KO.VP1-454AzK and 5F7-69-AzK were linked together using our established AAV-protein conjugation strategy described in the previous section (Figure 2d, S2). Upon mixing the site-specifically Tz-functionalized AAV and TCO-functionalized nanobody for 4 h, we observed the near-complete formation of the VP1-nanobody conjugate (KO.VP1-454-Nb) by SDS-PAGE analysis (Figure 2e). Next we assessed the transduction efficiency of WT-AAV2, KO.VP1-454-AzK, and KO.VP1-454-Nb against the HER2-overexpressing cell line SK-BR-3 (Figure 2f, S6a) at a constant MOI of 125. We also assessed their transduction efficiency against HEK293T cell line that does not overexpress the HER2 receptor. Each virus encoded a wild-type EGFP reporter gene, the expression of which was monitored via flow cytometry to quantify their transduction efficiency. As expected, KO.VP1-454AzK was non-infectious due to the detargeting mutations introduced at the HSPG-binding site. The attachment of nanobody to KO.VP1-454AzK (i.e., KO.VP1-454Nb) was found to dramatically enhance its infectivity toward HER2+ SK-BR-3 cells. Remarkably, KO.VP1-454Nb showed >8-fold improved transduction efficiency relative to WT-AAV2. By contrast, nanobody attachment onto KO.VP1-454AzK resulted in only a minor increase of its transduction efficiency toward HEK293T cells, which do not overexpress the HER2 receptor; the resulting virus KO.VP1-454Nb was much less infective relative to WT-AAV2 (Figure 2g, S6b). To further confirm that the impressive infectivity of KO.VP1-454Nb toward SK-BR-3 cells is indeed mediated by the target receptor (HER2), we showed that the addition of free nanobody inhibits the transduction efficiency of KO.VP1-454Nb in a dose-dependent manner (Figure 2h, S7). These experiments demonstrate that the attachment of antibody fragments provides an effective strategy to rationally reprogram the infectivity of AAV capsids toward new receptors.

### Optimization of the AAV-nanobody conjugate

A key strength of our approach is the high degree of control it offers on various aspects of the AAV-protein conjugation process, including: A) the site of attachment both on the virus and the protein; B) the number of proteins attached per capsid, by controlling the number of ncAAs introduced per capsid; and C) the length and chemical properties of the linker. Systematic optimization of these parameters has the potential to fine-tune the properties of AAV-protein conjugates for creating efficient gene-therapy vectors. To probe this hypothesis, we first explored the impact of the site of attachment on the performance of the AAV-nanobody conjugates. In addition to the original 454 site, we also introduced AzK at three additional surface-exposed sites within VP1 (Q263, T456, and R588), to generate KO.VP1-263AzK, KO.VP1-456AzK, and KO.VP1-588AzK (Figure 3a). The anti-HER2 5F7 Nb was successfully conjugated onto each AAV mutant, as demonstrated by the near-complete conversion of VP1 to a higher molecular weight band by SDS-PAGE analysis (Figure 3b). The transduction efficiency of each AAV2.KO mutant was evaluated toward SK-BR-3 cells before and after nanobody conjugation at a constant MOI. In each case, the KO virus was noninfectious, but nanobody attachment resulted in high levels of infectivity toward SK-BR-3 cells (Figure 3c, S8a), but not toward HEK293T cells (Figure S8b, S8c). The site of nanobody attachment was found to have a significant impact on transduction efficiency, with the KO.VP1-263AzK and KO.VP1-588AzK showing the lowest and highest activity, respectively. It is interesting to note that the optimal 588 residue is also part of the natural primary receptor binding site for AAV2.

**Figure 3:**
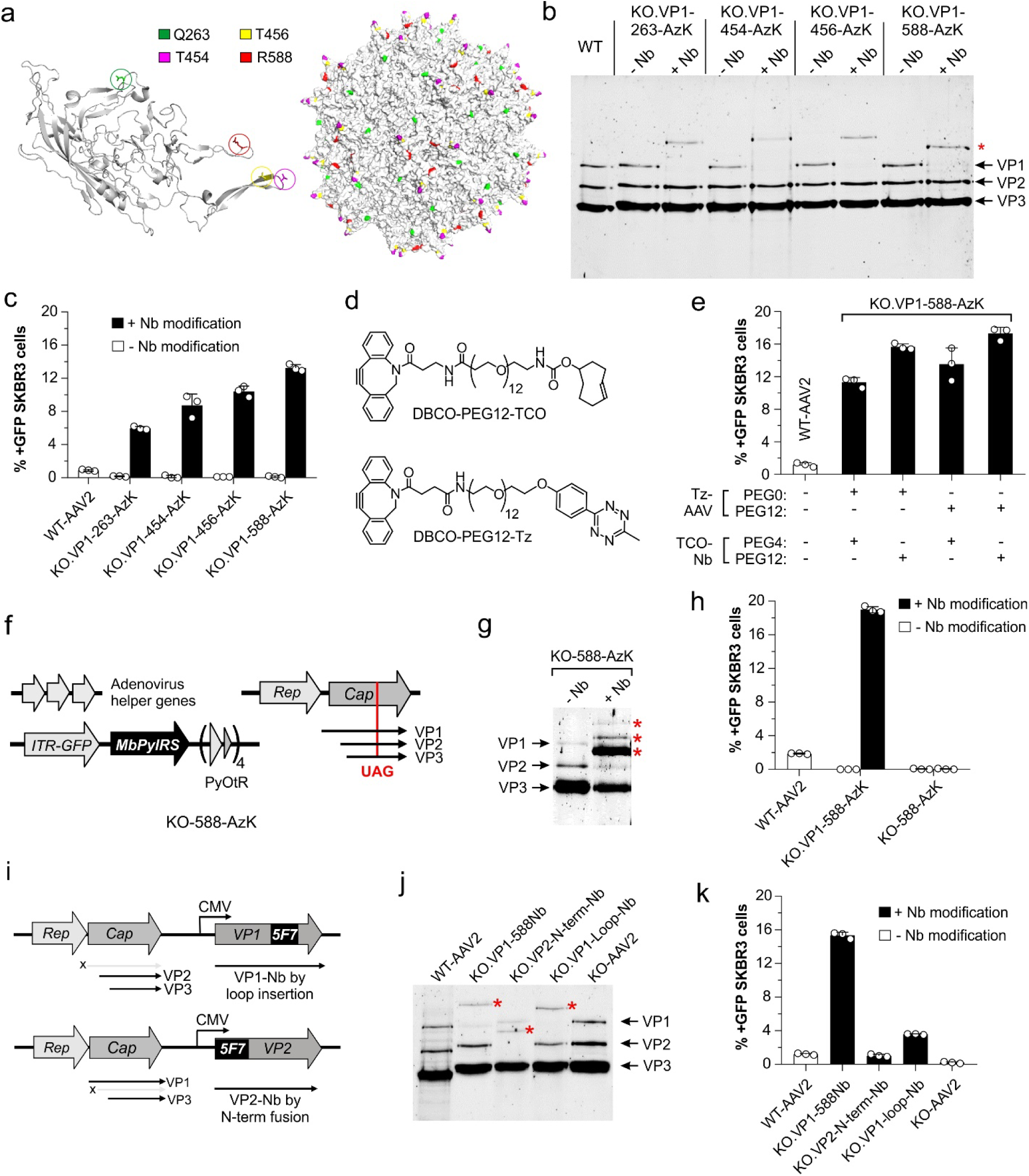
Optimization of the AAV2-nanobody conjugate. a) A color-coded depiction of the sites targeted for AzK incorporation in the AAV2 capsid. Left: Individual capsid protein. Right: The assembled capsid. b) SDS-PAGE analysis of AAV2-KO.VP1-Nb conjugate formation at four different sites on the VP1 protein. c) Infectivity of WT, unmodified and nanobody-conjugated KO.VP1-AzK mutants, measured by the percentage of GFP-expressing cells using flow cytometry, upon infecting SK-BR-3 cells at a constant MOI of 125. d) Structures of two longer bifunctional linkers used in this experiment. e) Infectivity of WT and KO.VP1-588Nb, generated using different combinations of bifunctional linkers with varying lengths. Infectivity was measured by percentage of GFP-expressing cells by flow cytometry, upon infecting SK-BR-3 cells at a constant MOI of 125. f) The plasmid system for packaging AAV2 site-specifically incorporating AzK at all 60 capsid proteins. g) SDS-PAGE analysis of the conjugation reaction between DBCO-PEG12-Tz-functionalized AAV2-KO-588-AzK and DBCO-PEG12-TCO-functionalized Nb-69-AzK. f) Infectivity of unmodified and nanobody-functionalized KO.VP1-588AzK (∼ 5 AzK per capsid) and KO-588-AzK (60 AzK per capsid). Infectivity was measured by the percentage of GFP-positive cells by flow cytometry, upon infecting SK-BR-3 cells at a constant MOI of 125. i) The plasmid system used to produce AAV2, where the nanobody is inserted into a permissible loop in VP1, or fused to the N-terminus of VP2. j) SDS-PAGE analysis of KO.VP1-588-Nb (chemical conjugate), KO.VP2-N-term-Nb, and KO.VP1-Loop-Nb. k) Infectivity of AAV2-WT, KO.VP1-588Nb (chemical conjugate), KO.VP2-N-term-Nb, KO.VP1-Loop-Nb, and KO-AAV2 measured by the percentage of GFP-positive cells by flow cytometry, upon infecting SK-BR-3 cells at a constant MOI of 125. All data shown as the mean ± s.d. of *n* = 3 replicates.

We also investigated the impact of the linker length between the AAV and Nb on retargeting efficiency. A combination of linkers with different lengths were tested: AAV mutant KO.VP1-588AzK, which showed the most efficient retargeting upon nanobody attachment, was labeled with either DBCO-Tz or DBCO-PEG12-Tz, while the Nb-69-AzK was labeled with either DBCO-PEG4-TCO or DBCO-PEG12-TCO (Figure 3d, S2). By linking the resulting Tz-functionalized AAV and TCO-functionalized 5F7 samples in different combinations, we generated a series of four AAV-nanobody conjugates with increasing linker length. All of these conjugates exhibited robust transduction efficiency toward SK-BR-3 cells (Figure 3e, S9). The linker-length was found to have a modest but significant impact; the conjugate with the longest linker showed roughly 50% higher transduction efficiency relative to the one with the shortest linker.

So far, we have generated the AAV-protein conjugates by introducing the ncAA handle only into the VP1 minor capsid protein, which is present in approximately 5 copies per capsid. We sought to probe how the attachment of a higher number of nanobodies on the virus capsid would affect the retargeting efficiency. To this end, we incorporated AzK at all 60 copies of the AAV.KO capsid, using previously described methods (Figure 3f). When we subjected the resulting virus to the conjugation protocol established above, successful nanobody conjugation to all three capsid proteins was observed by SDS-PAGE (Figure 3g). However, the conversion was incomplete (∼50%), likely due to excessive crowding on the capsid upon attaching a large number of proteins. Upon infecting SK-BR-3 cells, we observed a drastic decrease in the infectivity of this virus relative to KO.VP1-588Nb (Figure 3h, S10), highlighting the detrimental impact of over-modifying the capsid and the importance of controlled labeling.

Genetic approaches have also been used to display small antibody-like proteins on the AAV capsid.^4^ Such approaches include the fusion onto the minor capsid protein VP2, or insertion into a permissive loop on a minor capsid protein. We sought to compare the activity of our optimized AAV2-Nb chemical conjugate with such reported genetic fusion constructs between AAV2 and the same nanobody.^31–37^ Following the strategies reported by the work of Koch-Nolte^31^ and Buchholz et al.,^34^ we inserted the 5F7 nanobody sequence at the loop of the variable region IV of VP1, or fused it to the-N terminus of VP2 to generate constructs KO.VP1-Loop-Nb and KO.VP2-N-term-Nb, respectively (Figure 3i). SDS-PAGE analysis of the resulting virus preparations confirmed successful fusion/insertion of nanobody to the target minor capsid proteins (Figure 3j). Relative to AAV2.KO, which was non-infectious, KO.VP1-Loop-Nb and KO.VP2-N-term-Nb showed significant infectivity toward HER2+ SK-BR-3 cells, while maintaining low infectivity toward HEK293T cells (Figure 3k, S11). Gratifyingly, our AAV2-nanobody conjugate KO.VP1-588Nb significantly outperformed both of the genetic fusion constructs, showing approximately 4- and 15-fold higher infectivity relative to KO.VP1-Loop-Nb and KO.VP2-N-term-Nb, respectively (Figure 3k, S11). These experiments show that the unparalleled flexibility offered by our approach for linking together AAV2 and recombinant proteins enables the optimization of conjugates with superior retargeting efficiency.

### Synthesis of a full-length antibody-AAV conjugate

While some small and well-folded proteins such as nanobodies can also be displayed on the AAV capsid through genetic fusion, larger and more complex (e.g., multi-subunit) proteins are refractory to such strategies. By contrast, our method of generating protein-AAV conjugates can be theoretically extended to any recombinant protein that can be site-specifically chemically modified, either through ncAA mutagenesis or alternative strategies. To show that our approach can be used to generate functional conjugates between AAV and large and multi-subunit proteins, we aimed to attach the full-length anti-HER2 antibody Trastuzumab onto the AAV capsid. Given the broad and rich toolbox of antibodies available against countless targets, the ability to generate precise AAV-antibody conjugates is expected to be valuable for numerous applications. To this end, we first expressed full-length Trastuzumab incorporating AzK at site 122 in the light chain (LC) using the Expi293 expression system. Incorporation of AzK into the resulting antibody was confirmed by MS analysis (Figure S3). Then, to generate the AAV-Trastuzumab conjugate through IEDDA, KO.VP1-588AzK was labeled using DBCO-PEG12-Tz, and Trastuzumab-LC-122-AzK was labeled with DBCO-PEG12-TCO (Figure 4a, S3). Incubating Tz-labeled AAV2 and TCO-labeled Trastuzumab at room temperature for 4 hours resulted in efficient conjugate formation, as shown by the shift of the VP1 to a higher molecular-weight band in SDS-PAGE analysis (Figure 4b). Attachment of Trastuzumab onto non-infective KO.VP1-588AzK resulted in significantly higher infectivity relative to WT-AAV2 toward HER2+ SK-BR-3 cells (Figure 4c, S12a), but not HEK293T cells (Figure S12b-c), confirming successful retargeting. The ability to attach a full-length antibody – a 150 kDa protein with four subunits – onto the AAV2 capsid showcases the generality of our approach, which can be further exploited to generate a diverse class of AAV-protein conjugates to further engineer its properties.

**Figure 4:**
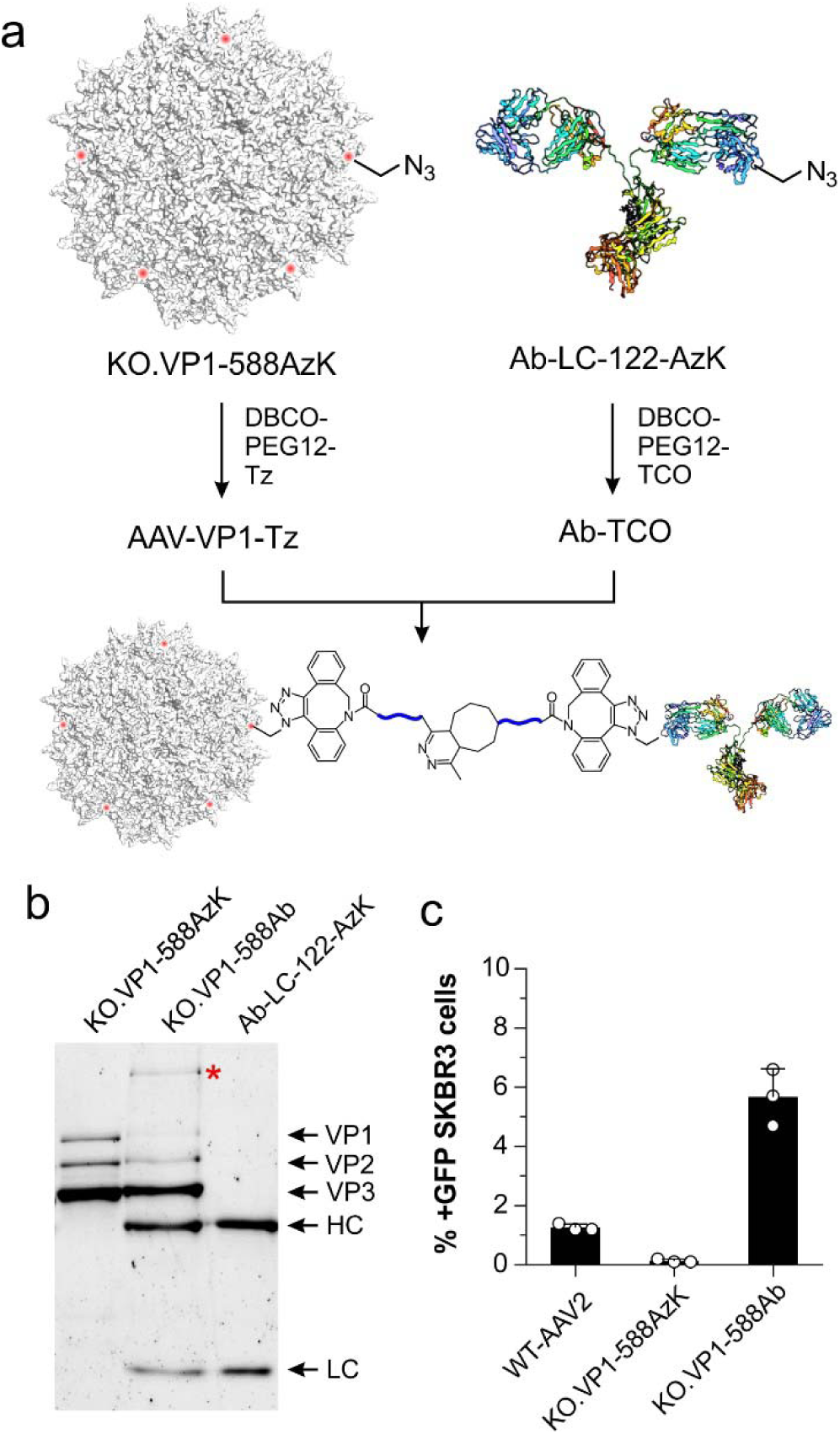
Attachment of a full-length antibody onto AAV. a) The scheme used to conjugate Trastuzumab-LC-122-AzK with AAV2-KO.VP1-588AzK. Virus-antibody conjugation was performed by incubating approximately 1.0 x 10^9^ gc/μL of AAV-VP1-Tz and 0.5 μM Ab-TCO at RT for 4 hours. b) SDS-PAGE (reducing) analysis of the conjugation reaction between AAV-VP1-Tz and Ab-LC-TCO. The asterisk highlights the band corresponding to the expected conjugate between VP1 and Trastuzumb-LC. c) Infectivity of WT-AAV2, unmodified and Ab-modified KO.VP1-588AzK, measured by the percentage of GFP-positive cells by flow cytometry, upon infecting SK-BR-3 cells at a constant MOI of 250. Data shown as the mean ± s.d. of *n* = 3 replicates.

### Efficient and selective gene delivery by AAV2-nanobody conjugates *in vivo*

In AAV gene therapy, a key goal is to create capsids that selectively deliver genetic cargo to specific tissues upon systemic administration. However, existing AAV vectors often exhibit off-target infectivity, especially toward the liver. Encouraged by the efficient and selective gene delivery exhibited by our optimized AAV2-5F7 conjugates toward HER2+ cancer cells *in vitro*, we sought to characterize its efficacy *in vivo* in a mouse model. To this end, we first characterized the transduction efficiency of KO.VP1-588Nb toward BT474 cells, another HER2-overexpressing cancer cell line that enables more facile xenograft formation. Evaluating the transduction efficiencies toward BT474 cells revealed the same infectivity pattern observed for SK-BR-3: KO.VP1-588AzK was non-infectious but attachment of 5F7 to generate KO.VP1-588Nb resulted in robust transduction of these cells, at a significantly higher level relative to WT-AAV2 (Figure 5a, S13).

**Figure 5:**
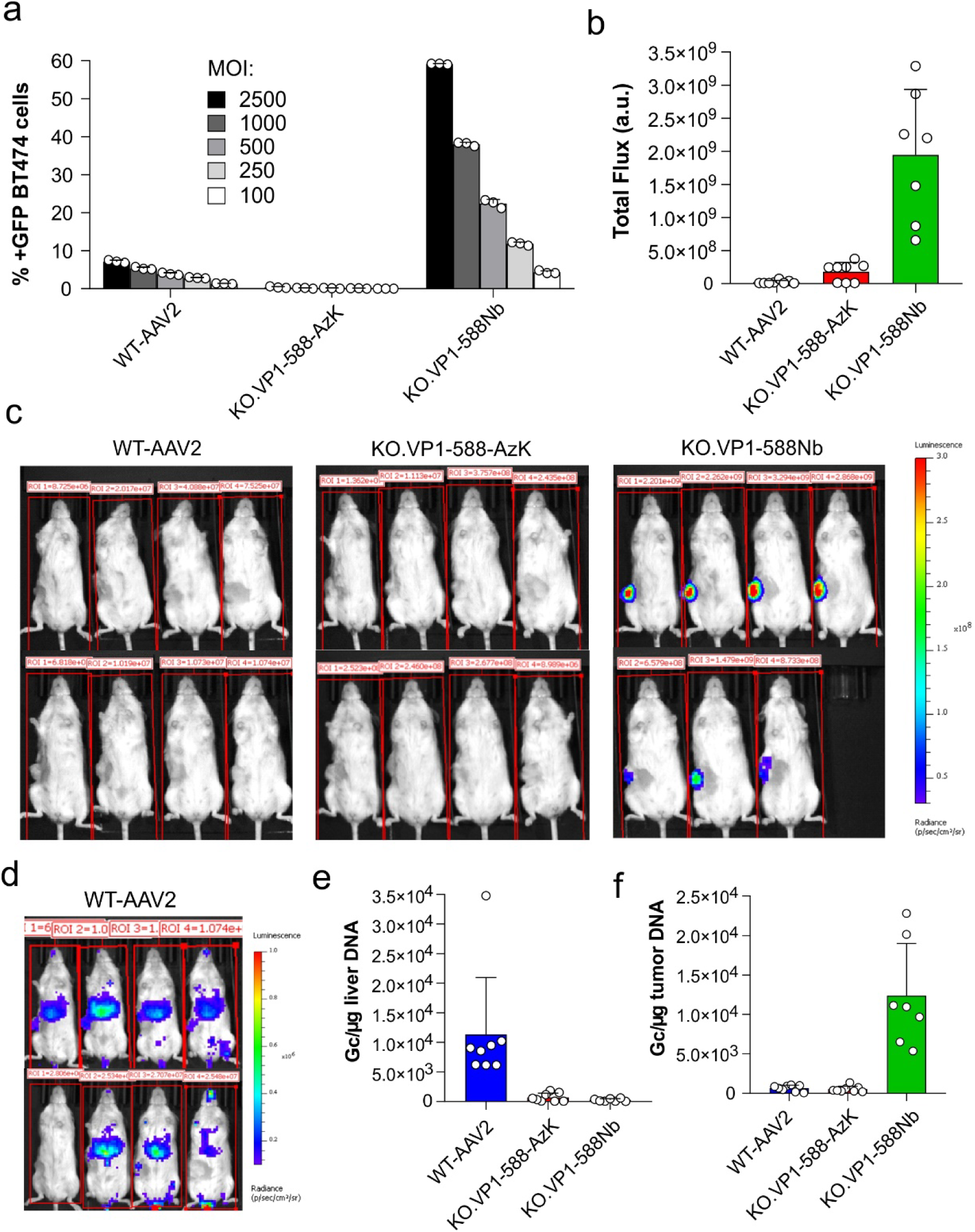
AAV2-Nb conjugate enables selective and efficient transduction of HER2+ tumor cells *in vivo*. a) *In vitro* infectivity of AAV2-WT, unmodified and Nb-modified KO.VP1-588-AzK, measured by the percentage of GFP-expressing cells by flow cytometry, upon infecting BT474 cells at following MOIs: 100, 250, 500, 1000, 2500. Data shown as the mean ± s.d. of *n* = 3 replicates. b) Signal from luminescence imaging of mice (harboring a BT474 tumor xenograft) injected with WT-AAV2, KO.VP1-588-AzK, and KO.VP1-588-Nb, 14 days post-injection. Eight mice per group were intravenously injected with 1.0 x 10^11^ gc of each virus. c) Whole-body bioluminescence images (ventral) of mice associated with the experiment in panel b, 14 days post-injection (fixed luminescence scale). d) Magnified ventral images of the 8 mice injected with WT-AAV2 showed in an enhanced luminescence scale to visualize the weak signal, which originates largely from the liver. AAV-encoded transgene in DNA retrieved from liver (e) and tumor (f) tissues, measured by qPCR, harvested at the end of the *in vivo* study (5 weeks post-injection). Data shown as the mean ± s.d. of *n* = 8 mice for WT, KO.VP1-588-AzK viruses and n=7 mice for KO.VP1-588Nb (one mouse expired on day 12 with no obvious cause of death).

Next, three groups of mice (n=8) were injected with 5.00 x 10^6^ BT474 cells/mouse for xenograft formation, and tumor volume was allowed to expand to approximately 100 mm^3^. Identical doses of WT, KO.VP1-588AzK and KO.VP1-588-Nb viruses encoding a luciferase reporter gene were delivered to each group of mice by tail-vein injection. Whole-body luminescence imaging of the animals was performed on a weekly basis starting from the 2^nd^ week (Figure 5c, S14). After 5 weeks, the animals were sacrificed and tumors and livers were harvested for transgene titering. As expected, luminescence imaging of mice infected with WT-AAV2 on the 2^nd^ week revealed low levels of luminescence originating largely from the liver (Figure 5c-d). By contrast, KO.VP1-588-Nb exhibited a massively higher levels of luminescence (>85 fold), and the signal originated selectively from the tumor. Not surprisingly, as the tumor rapidly expanded in the subsequent weeks, the luminescence levels from tumor reduced due to the dilution of the delivered transgene (Figure S14). Nonetheless, qPCR analysis of the tumor samples harvested after 5 weeks still showed >19-fold more efficient transgene delivery into the tumor by KO.VP1-588-Nb relative to WT-AAV2 (Figure 5f), further corroborating the imaging experiments. By contrast, qPCR analysis showed severely attenuated (>45-fold lower) transgene delivery into the liver by KO.VP1-588Nb relative to WT-AAV2 (Figure 5e). These results show that the chemical conjugation of antibody fragments can remarkably improve both the efficiency and selectivity of AAV-mediated gene delivery *in vivo*. While KO.VP1-588-AzK exhibits limited infectivity *in vitro* in cell culture models, consistent with prior findings, it does demonstrate some infectivity *in vivo*.^65^

## Conclusions

The ability to display proteins such as antibody fragments onto the AAV capsid has been explored as an attractive avenue to rationally engineer the properties of this promising gene therapy vector. However, due to the delicate nature of the intricate AAV capsid, traditional genetic approaches have been severe restricted in their scope; such approaches are only compatible with small and well-folded proteins, and there are only a limited number of ways to graft them onto the capsid proteins. Here, we report an alternative chemical strategy to synthesize AAV-protein conjugates that offer a remarkable degree of versatility, allowing systematic optimization of the conjugate. Our strategy uses GCE to efficiently introduce a ncAA into both AAV and the recombinant protein in a site-specific manner, and then links them together using synthetic bispecific linkers by ultrafast IEDDA bioorthogonal conjugation chemistry. This strategy was used to seamlessly attach three different proteins, sfGFP, 5F7 nanobody, and a full-length antibody Trastuzumab, onto the minor capsid protein VP1 of AAV2. Using this method, it was possible to systematically alter the site of attachment, the number of proteins attached per capsid, and the properties of the linker, which enabled us to optimize an AAV2-5F7 nanobody conjugate that exhibits highly efficient and selective transduction of HER2+ cancer cells. In mouse models, the optimized AAV2-5F7 nanobody conjugate facilitated remarkably efficient transgene delivery into a HER2+ tumor xenograft, while exhibiting negligible off-target delivery. The same strategy also enabled the synthesis of a functional AAV2-Trastuzumab (full-length antibody) conjugate, highlighting the generality of this approach. It should be possible to readily adapt our workflow to retarget AAV to additional cell-surface receptors, by attaching a suitable antibody or antibody-like proteins. Given the art of developing antibodies against desired cell-surface receptors is well-established, this provides a highly modular and rational avenue to generate efficient and selective gene therapy vectors. Finally, attachment of recombinant proteins onto the AAV capsid can also enable rational engineering of other therapeutically relevant properties such as immunogenicity and pharmacokinetics.

## ASSOCIATED CONTENT

### Supporting Information

Experimental methods, nucleotide sequences, supplementary figures and tables. (PDF)

## AUTHOR INFORMATION

**Notes**: A patent application has been submitted based on the technology described herein, where AC and QP are co-inventors. AC is a cofounder and senior advisor at BrickBio, Inc.

## Acknowledgement

AC acknowledges support from NIH (R35GM136437) and National Science Foundation (2128185)

