## Supporting information for "A facile chemical strategy to synthesize precise AAV-protein conjugates for targeted gene delivery"

### Selective viral transduction of AAV via control chemical conjugation with Antibody

**Extended data**

Unprocessed LC-MS of sfGFP-151-AzK **1**.

Calculated: 27675; found: 27673

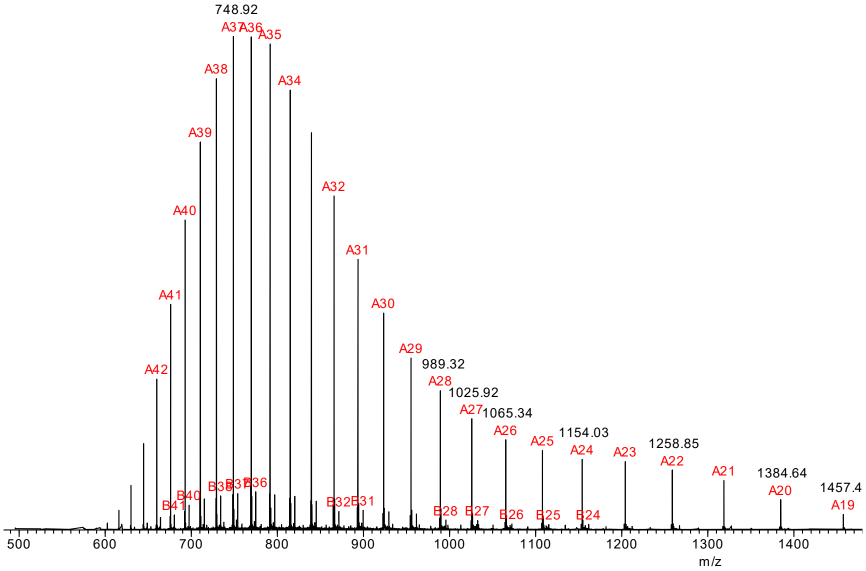


Unprocessed LC-MS of sfGFP-151-AzK and DBCO-PEG4-TCO adduct **4**

Calculated: 28351; found: 28349

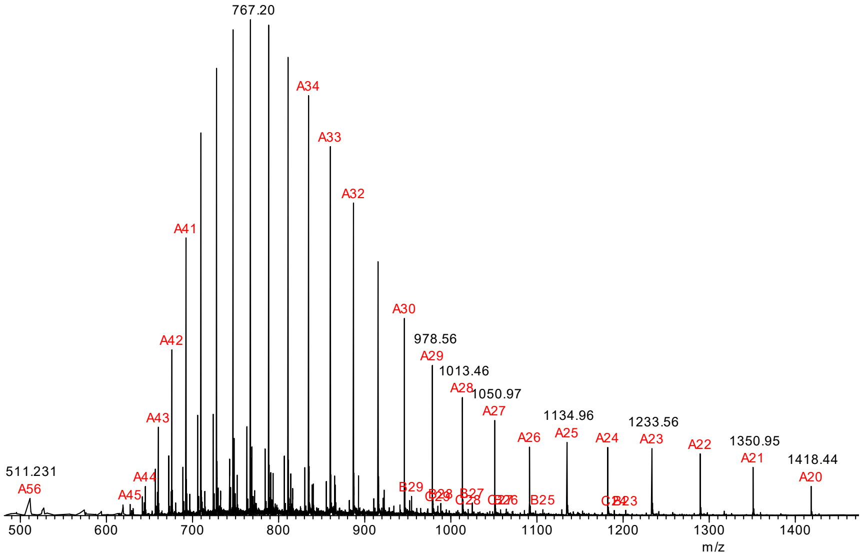

Unprocessed LC-MS of sfGFP-151-AzK and DBCO-sulfo-Tz adduct **5.**

Calculated: 28359; found: 28358

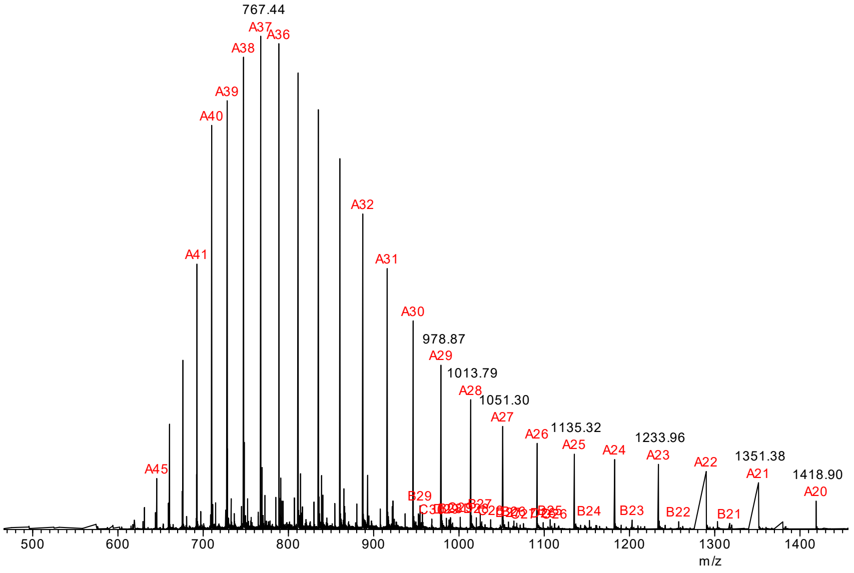


Unprocessed LC-MS of sfGFP coupling product **6**

Calculated: 56682; found: 56675

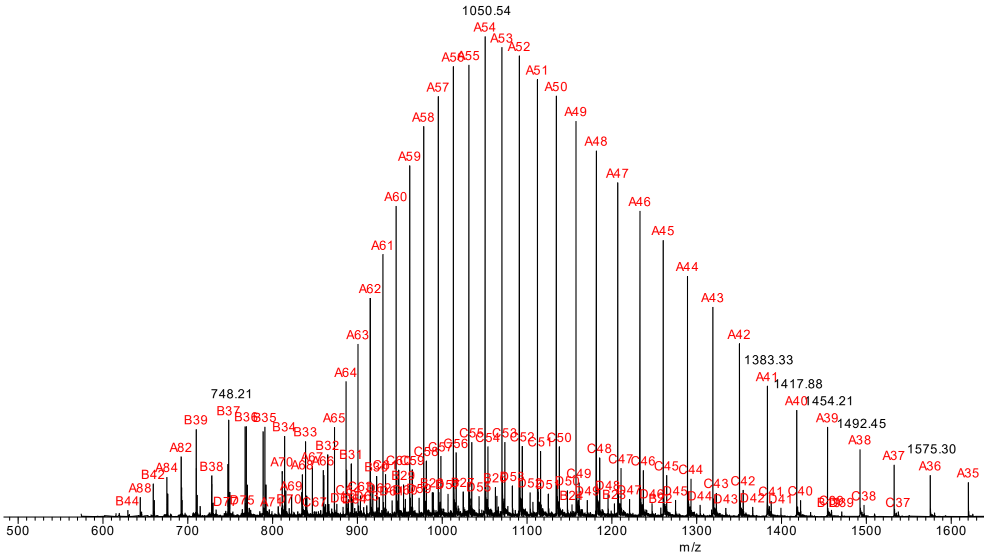


**Figure S1**: Full ESI-MS spectra for the experiments shown in Figure 1e.

Processed and unprocessed (below) LC-MS of AntiHer2-Nb-69-AzK

Calculated: 13910; found: 13910

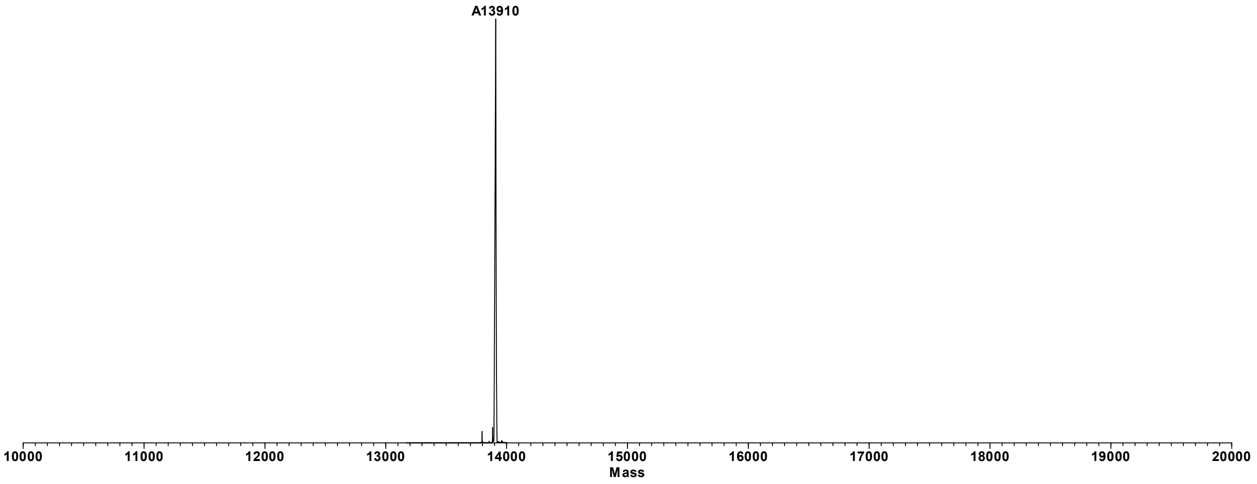


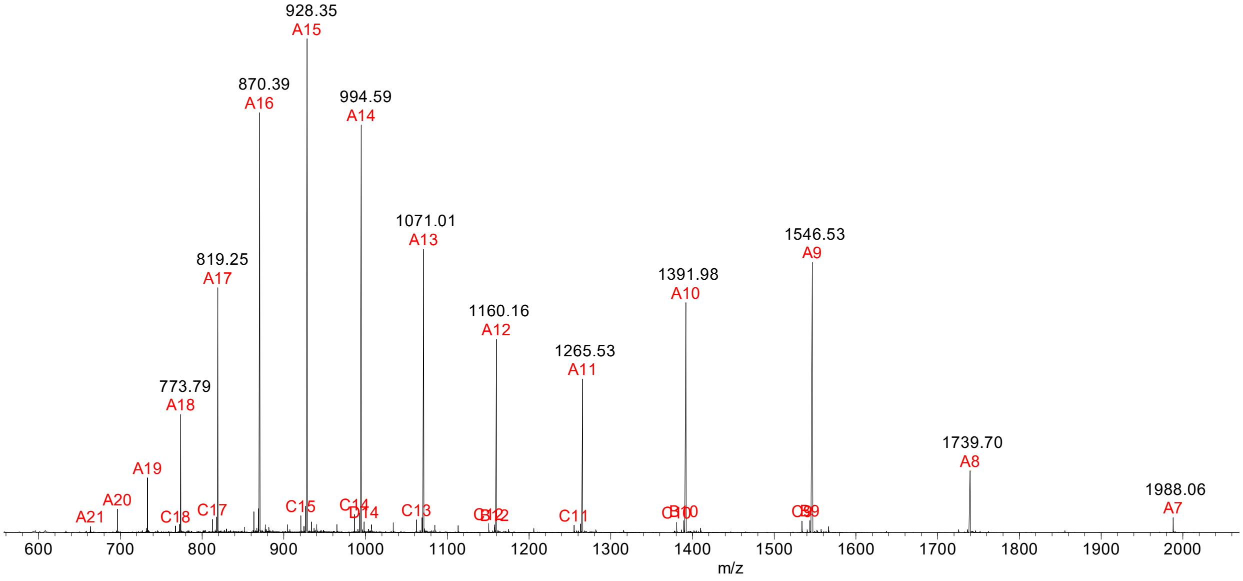

Processed and unprocessed (below) LC-MS of AntiHer2-Nb-69-AzK and DBCO-PEG4-TCO adduct

Calculated: 14585; found: 14585

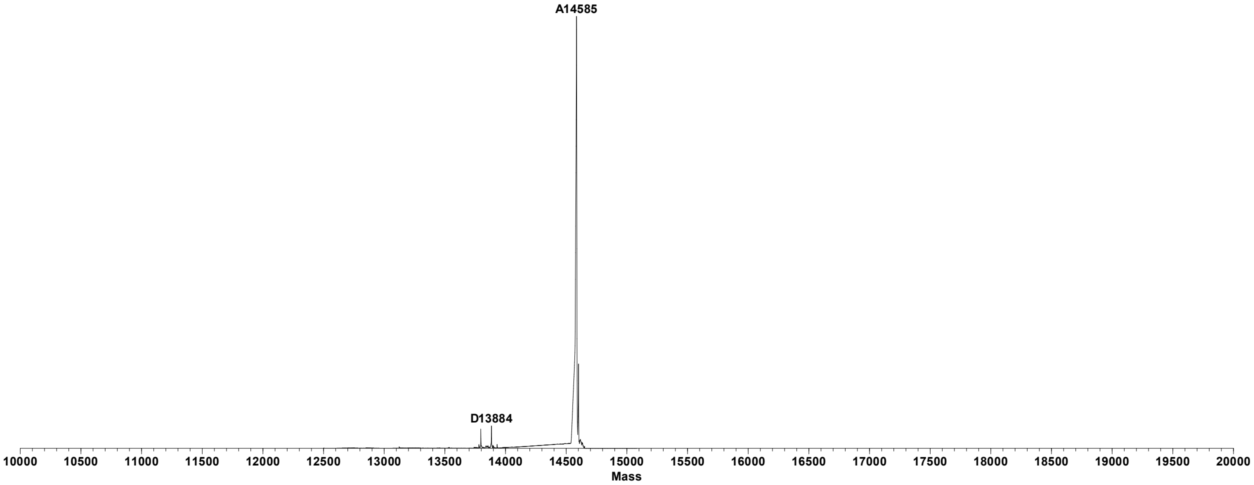


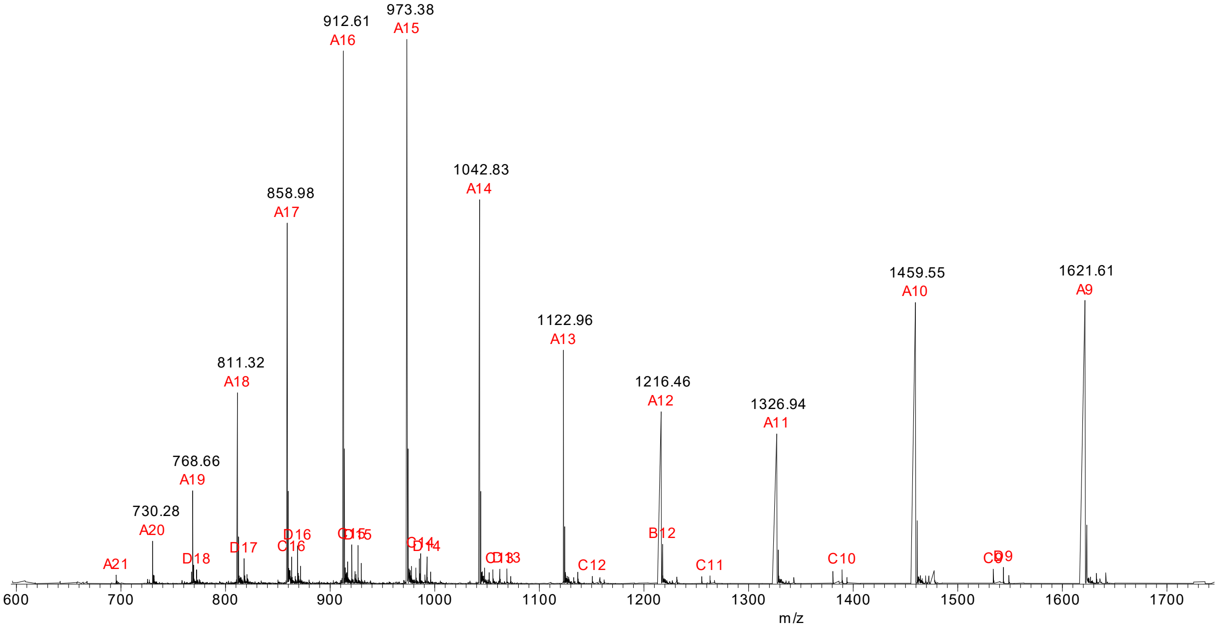

Processed and unprocessed (below) LC-MS of AntiHer2-Nb-69-AzK and DBCO-PEG12-TCO adduct

Calculated: 14938; found: 14938

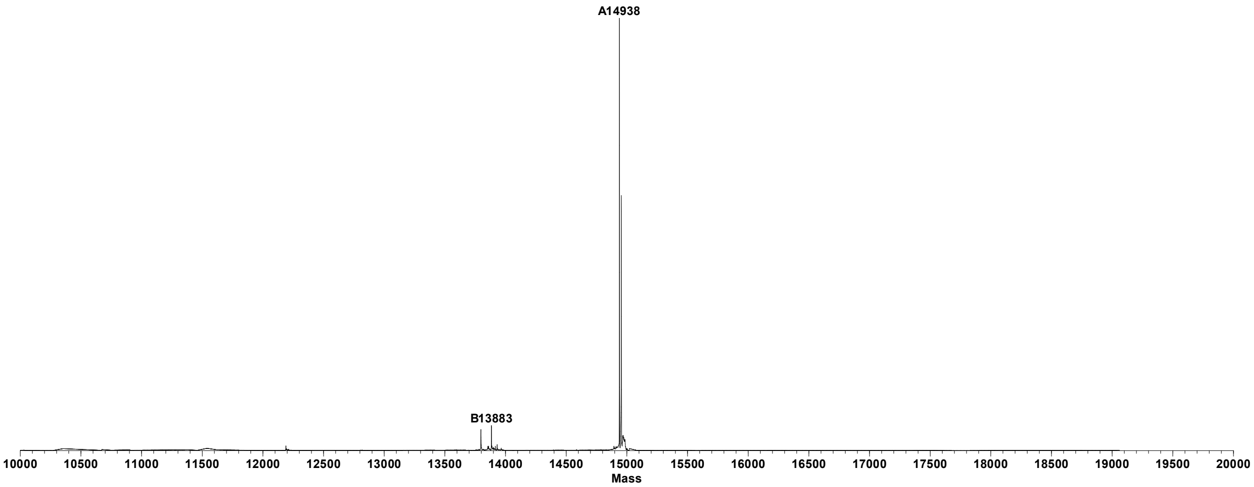


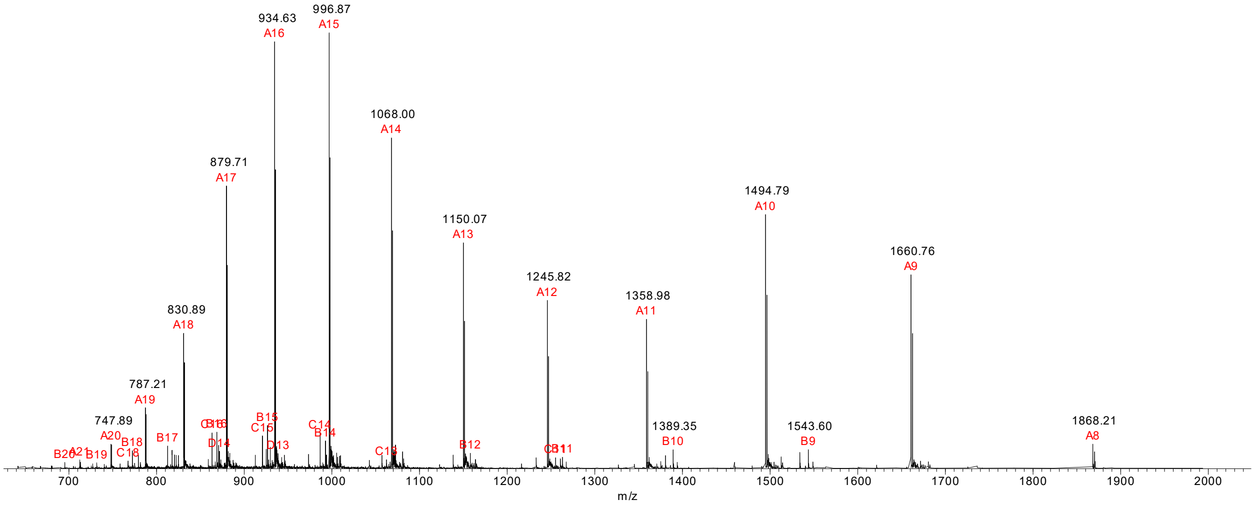


**Figure S2**: Deconvoluted and full ESI-MS spectra for the conjugation of Anti-Her2-Nanobody with DBCO-PEG4-TCO shown in Figure 2d and with DBCO-PEG12-TCO shown in Figure 3e.

Processed and unprocessed (below) LC-MS of Trastuzumab-LC-122-AzK

Light chain: Calculated: 23569; found: 23566

Heavy chain: Calculated: 49211; found: 49208

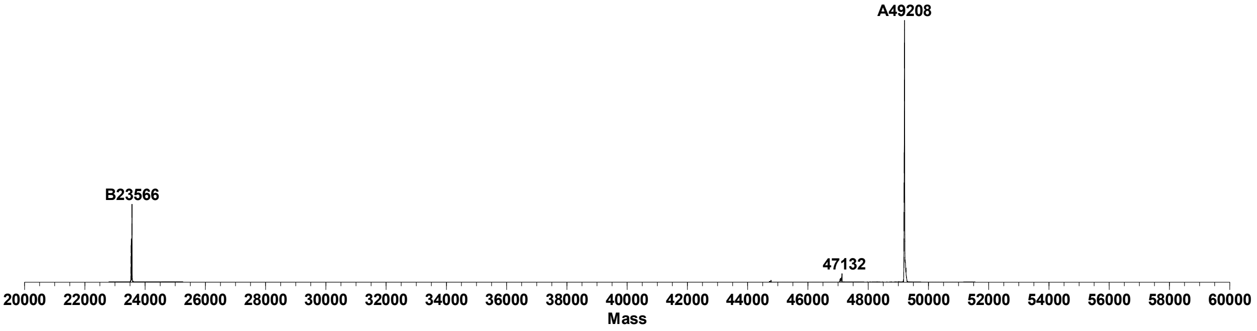


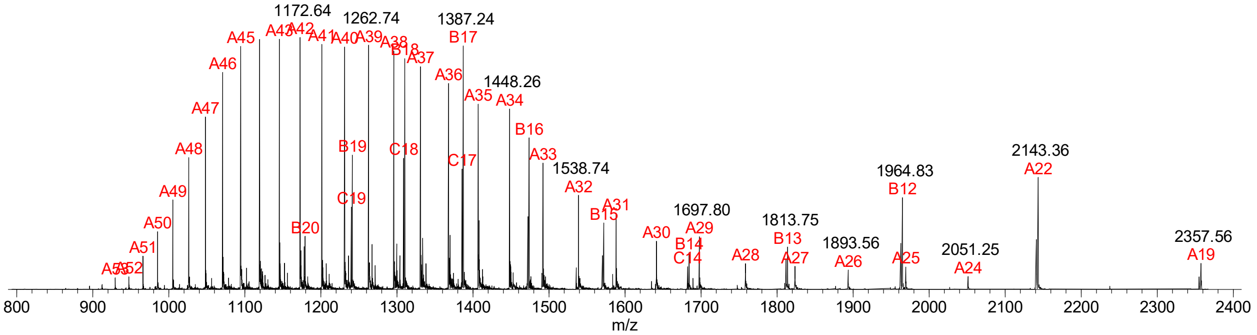

Processed and unprocessed (below) LC-MS of Trastuzumab-LC-122-AzK and DBCO-PEG12-TCO adduct, followed by addition of Tetrazine-aniline

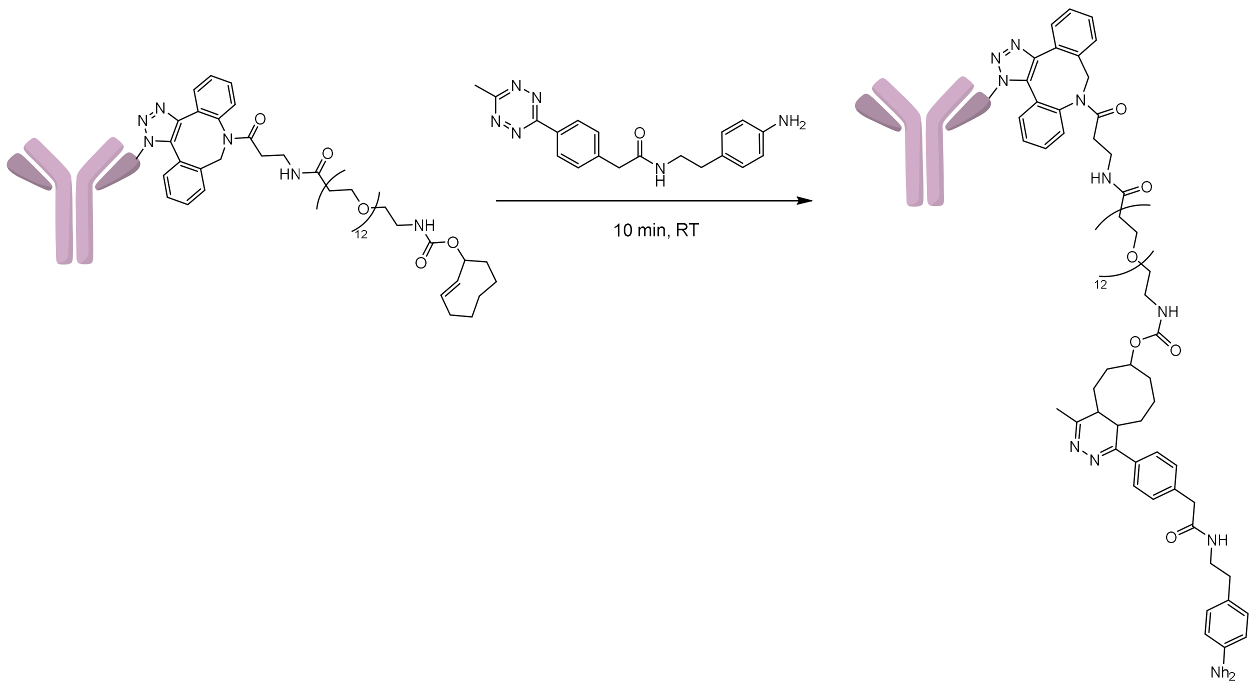


Light chain: Calculated: 24917; found: 24915
Heavy chain: Calculated: 49211; found: 49213

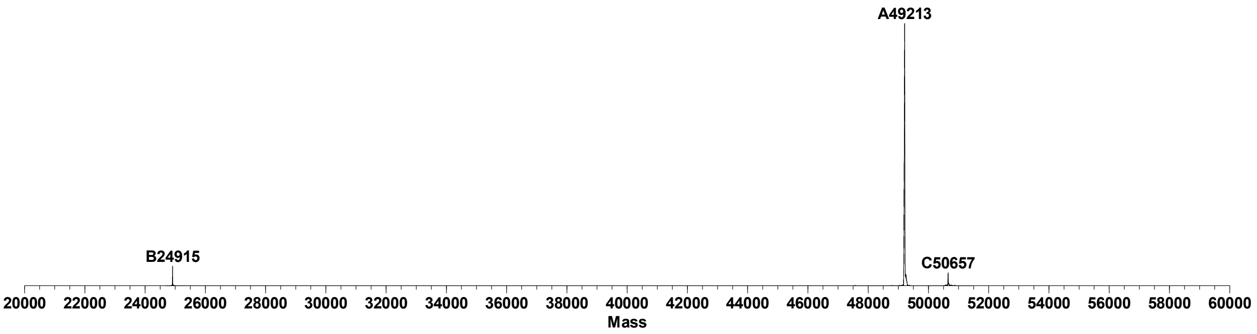


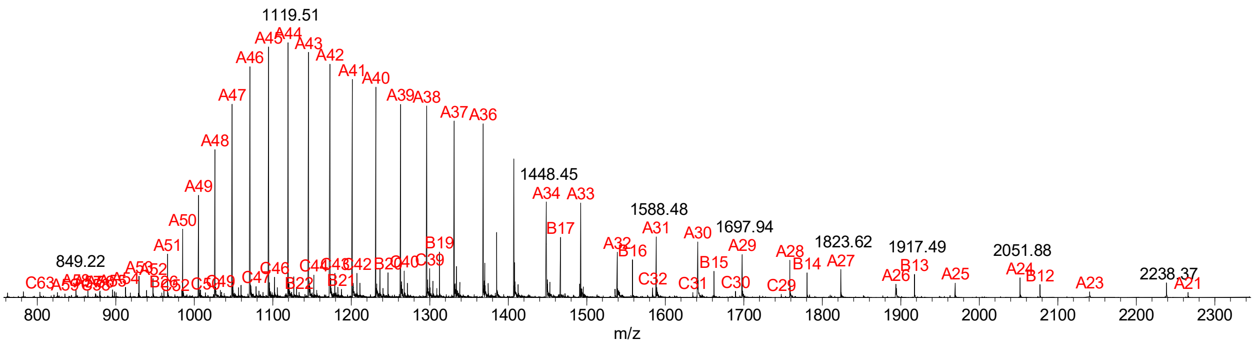

Zoomed-in processed LC-MS of the light chain

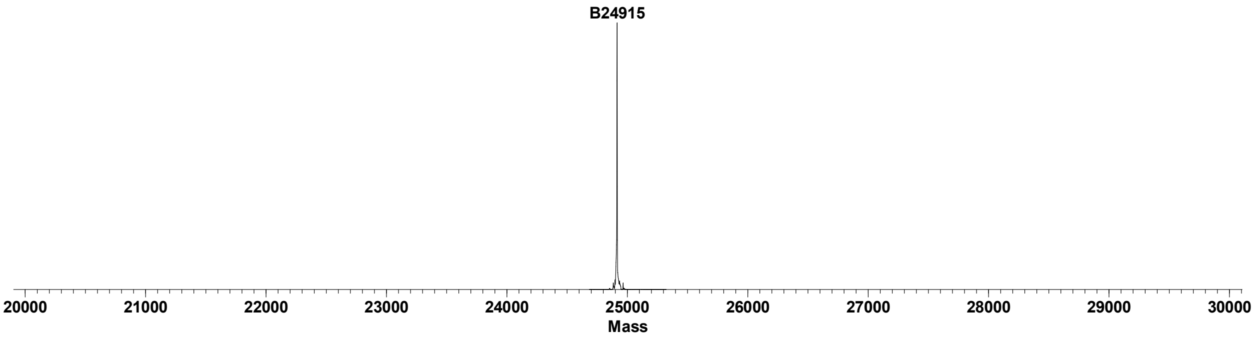


**Figure S3**: Deconvoluted and full ESI-MS spectra for the conjugation of Anti-Her2-Antibody-LC-122-AzK and DBCO-PEG12-TCO adduct shown in Figure 4, followed by addition of Tetrazine-aniline for the protection of TCO group during antibody treatment for LC-MS.

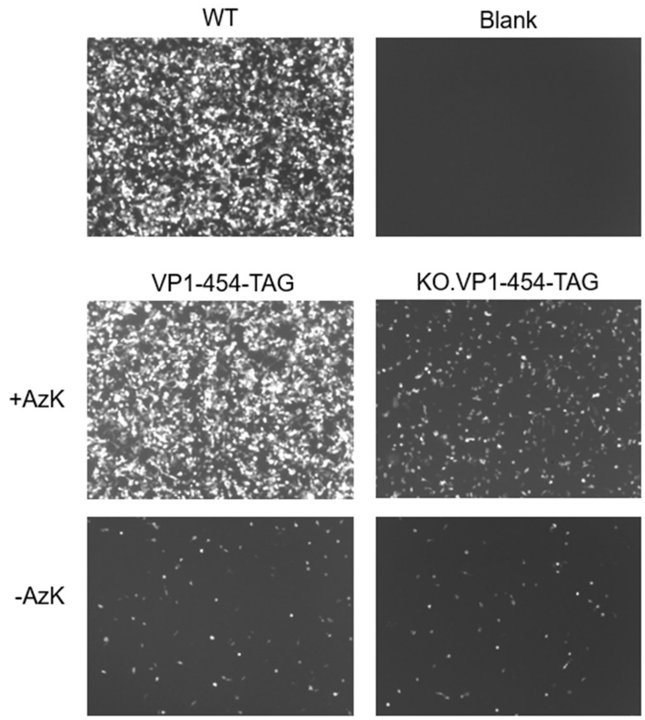


**Figure S4**: Representative fluorescence images of cells associated with the experiment described in Figure 2c. HEK293T cells were infected with a constant MOI (50) of AAV2 wild-type (WT) or AzK mutant at VP1 protein without or with detargeting from heparan sulfate by R585A and R588A mutations (KO), and imaged 48 h post-infection.

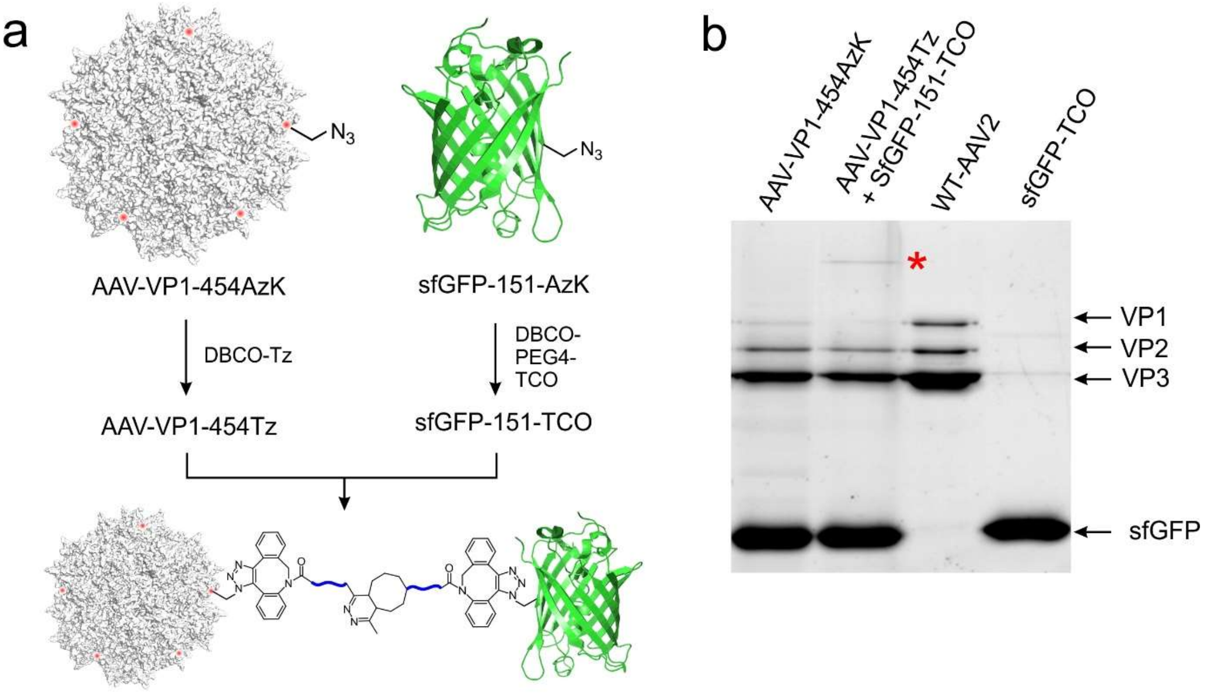


**Figure S5**. a) The scheme used to chemically conjugate sfGFP-151-AzK and AAV2-VP1-454-AzK. b) SDS-PAGE analysis of the conjugation reaction between AAV-VP1-454Tz and sfGFP-151-TCO. The asterisk highlights the band corresponding to the conjugate between VP1 and sfGFP.

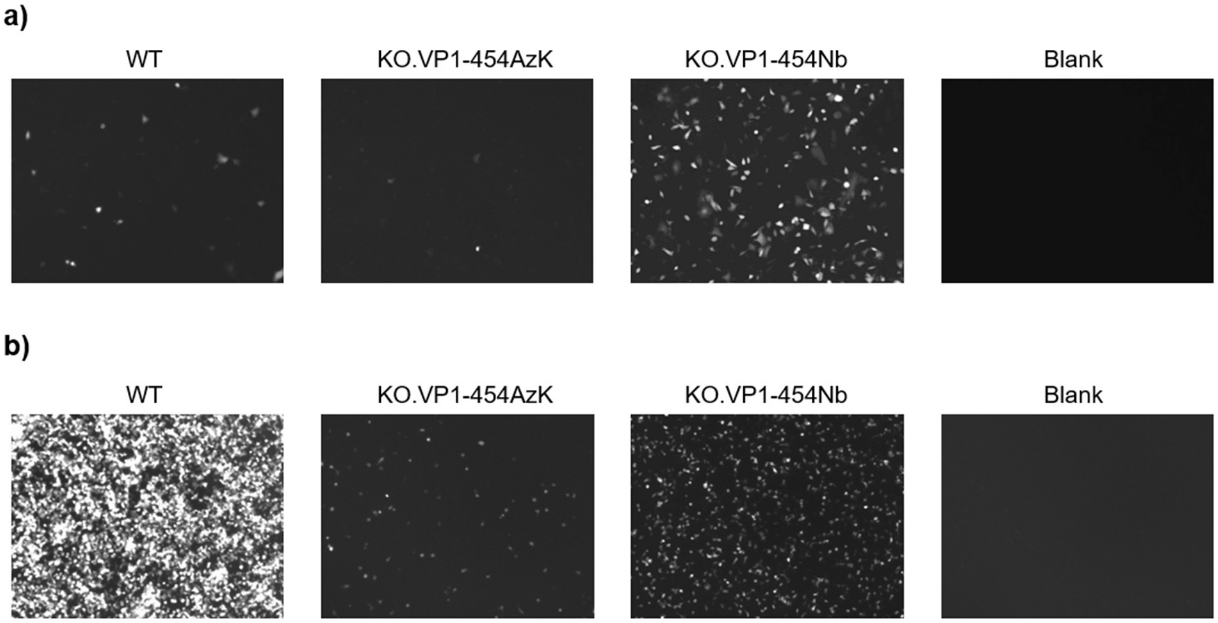


**Figure S6**: Representative fluorescence images of cells associated with the experiment described in Figure 2f and 2g. a) SK-BR-3 cells were infected with a constant MOI 125 of AAV2 WT or detargeted virus without (KO.VP1-454AzK) or with conjugation with AntiHer2-Nanobody (KO.VP1-454Nb), and imaged 48 h post-infection. b) HEK293T cells were infected with a constant MOI 50 of WT-AAV2 or detargeted virus without (KO.VP1-454AzK) or with conjugation with AntiHer2-Nanobody (KO.VP1-454Nb), and imaged 48 h post-infection.

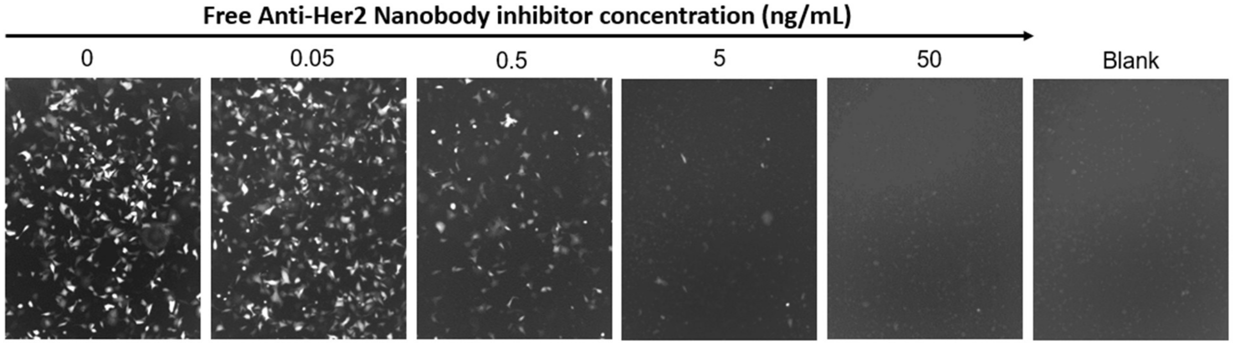


**Figure S7**: Fluorescence images of cells associated with the experiment described in Figure 2h. SK-BR-3 cells were infected with a constant MOI 500 of KO.VP1-454Nb with increasing concentration of free AntiHer2-Nanobody added as an inhibitor, and imaged 48 h post-infection.

a)

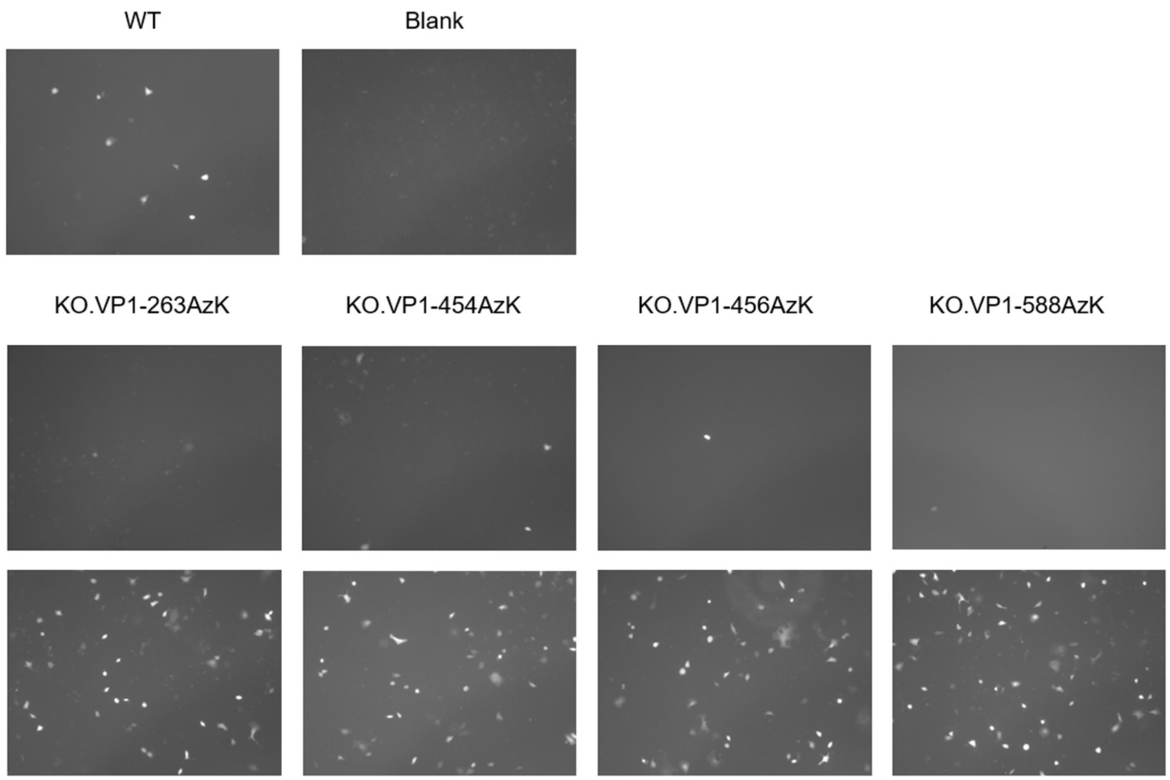


b)

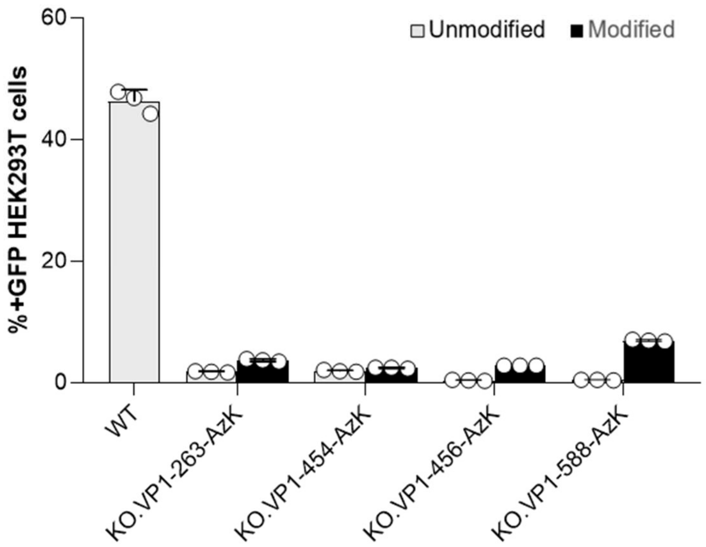


(continued on the next page)

c)

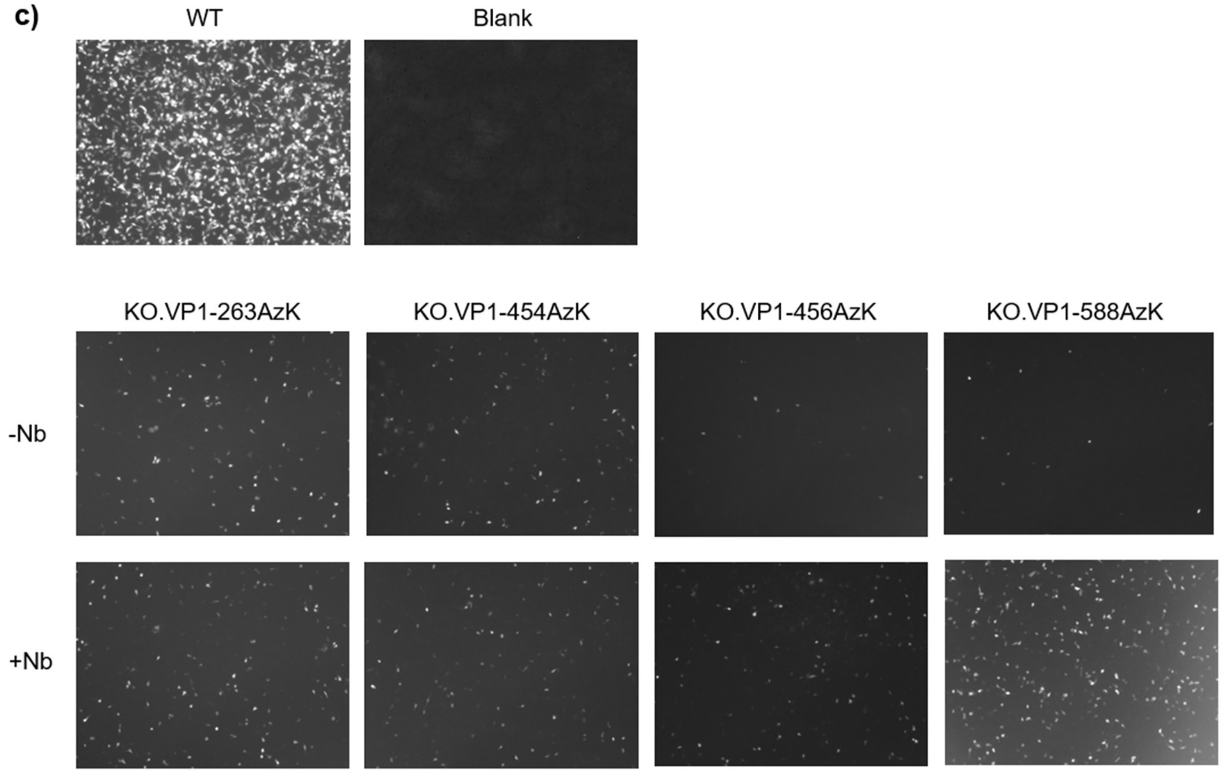


**Figure S8**: a) Representative fluorescence images of cells associated with the experiment described in Figure 3c. SK-BR-3 cells were infected with a constant MOI 125 of AAV2 WT, unmodified KO.VP1-AzK site-variants, and their modified counterparts with AntiHer2-Nanobody, and imaged 48 h post-infection. b) Infectivity of WT, unmodified KO.VP1-AzK site-variants, and their modified counterparts with AntiHer2-Nanobody, measured by the percentage of positive GFP cells by flow cytometry, upon infecting HEK293T cells at a constant MOI 50. Data given as the mean ± s.d. of $n = 3$ replicates. c) Fluorescence images associated with panel b.

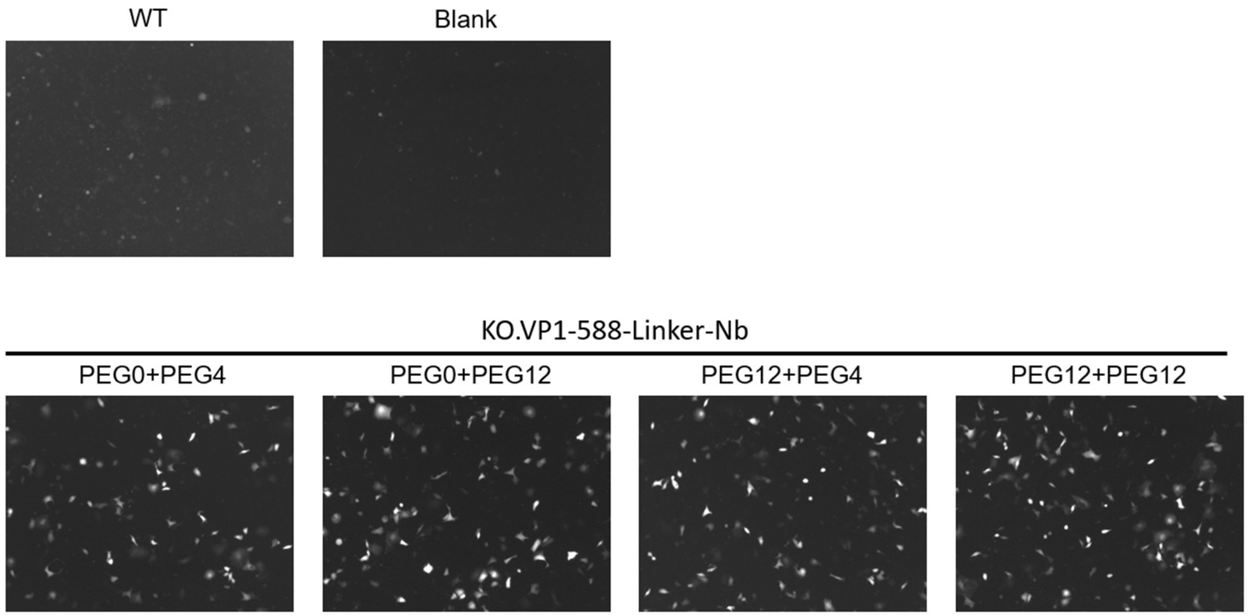


**Figure S9**: Representative fluorescence images of cells associated with the experiment described in Figure 3e. SK-BR-3 cells were infected with a constant MOI 125 of AAV2 WT, AntiHer2-Nanobody conjugated viruses at site 588 of VP1 protein with different linker combinations, and imaged 48 h post-infection.

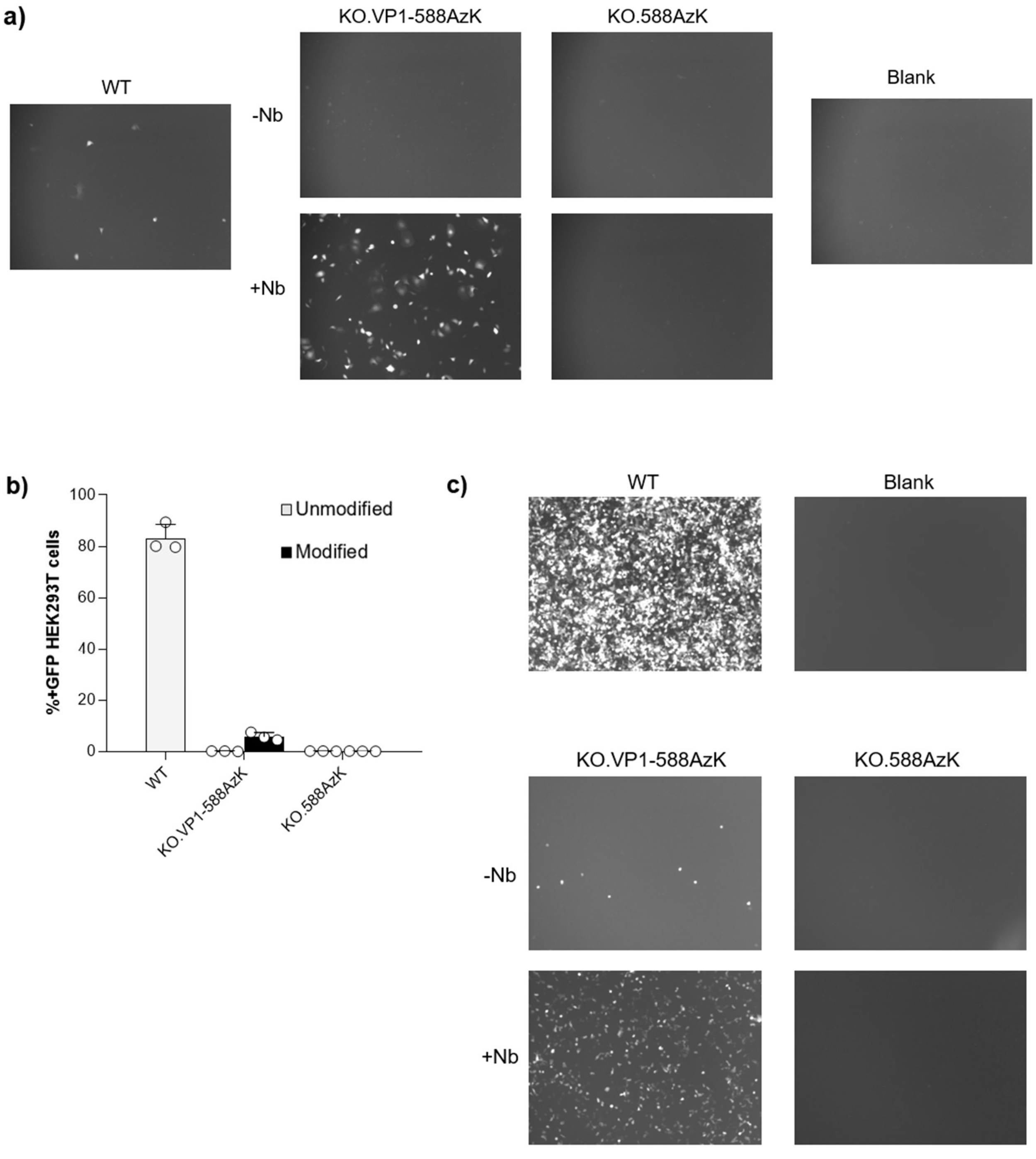


**Figure S10**: a) Representative fluorescence images of cells associated with the experiment described in Figure 3h. SK-BR-3 cells were infected with a constant MOI 125 of AAV2 WT, AntiHer2-Nanobody conjugated viruses with different stoichiometry at site 588 of VP1 or VP1+2+3 proteins, and imaged 48 h post-infection. b) Infectivity of WT, unmodified and modified virus at site 588 of VP1 or VP1+2+3 proteins with AntiHer2-Nanobody, measured by the percentage of positive GFP cells by flow cytometry, upon infecting HEK293T cells at a constant MOI 50. Data given as the mean ± s.d. of $n = 3$ replicates.  c) Fluorescence images associated with panel b.

**a)**

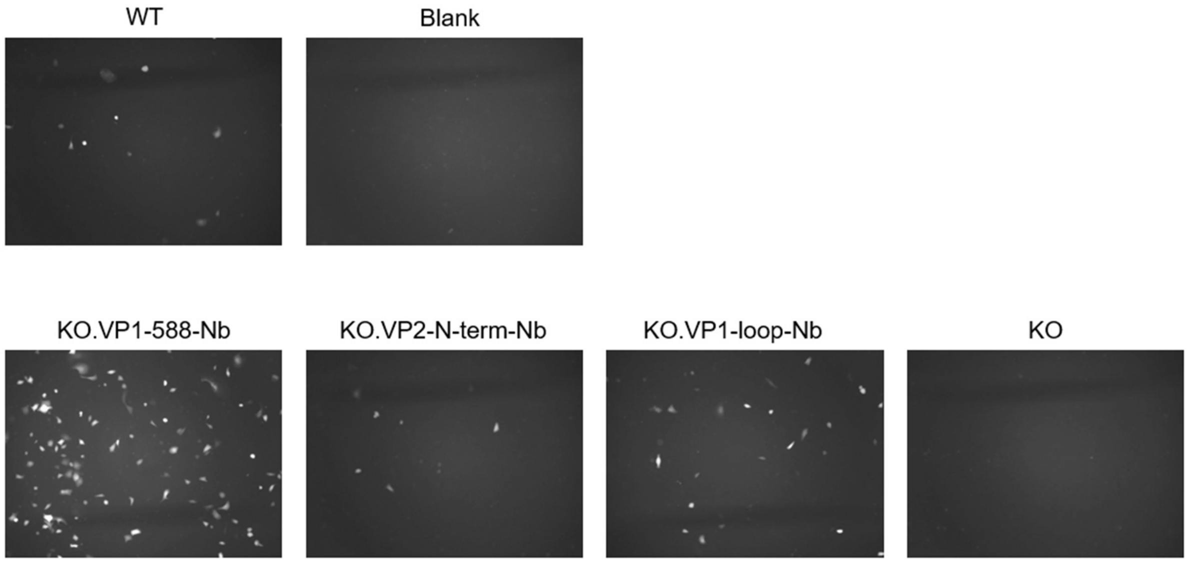


**b)**

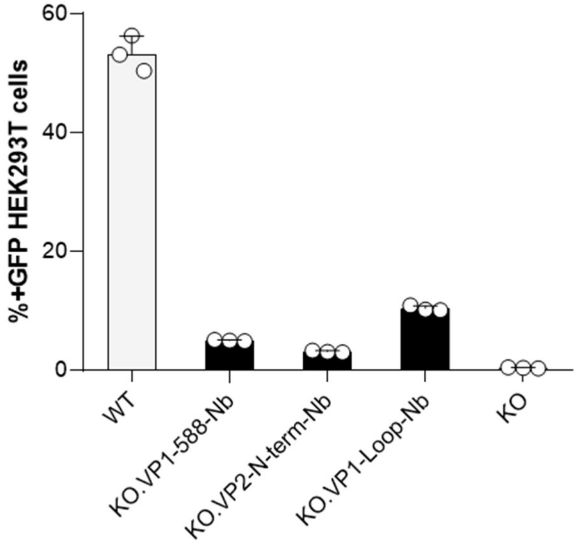


(continued on the next page)

**c)**

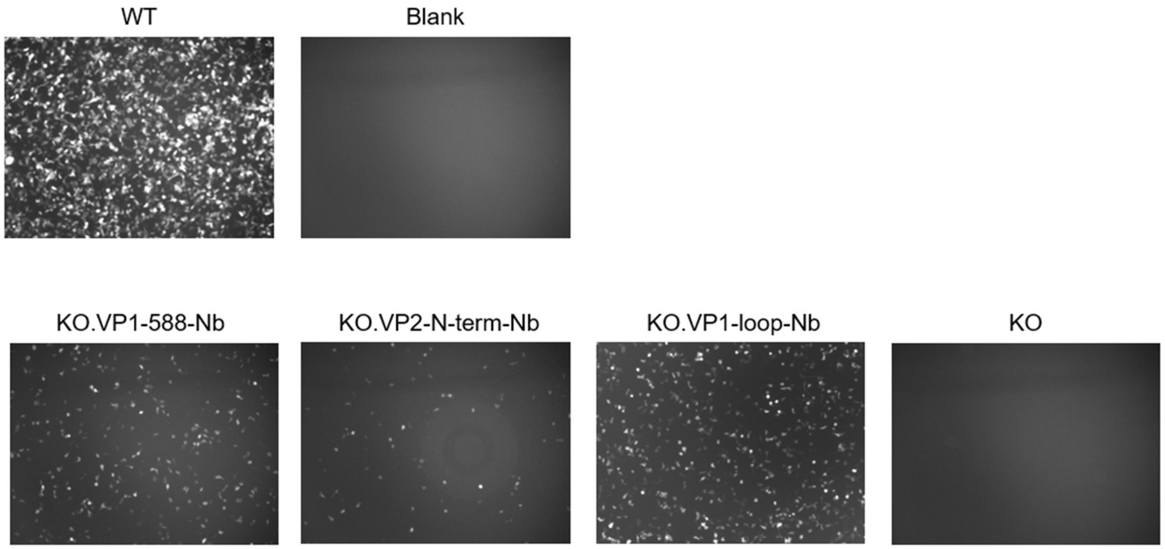


**Figure S11**: a) Representative fluorescence images of cells associated with the experiment described in Figure 3k. SK-BR-3 cells were infected with a constant MOI 125 of AAV2 WT, AntiHer2-Nanobody chemically conjugated virus at site 588 (KO.VP1-588Nb), virus with AntiHer2-Nanobody genetically fused to N terminus of VP2 protein virus (KO.VP2-N-term-Nb), Virus with AntiHer2-Nanobody genetically inserted to the loop of VP1 protein (KO.VP1-loop-Nb), and detargeted virus by R585A, R588A mutations (KO), and imaged 48 h post-infection. b) Infectivity of AAV2-WT, KO.VP1-588Nb, KO.VP2-N-term-Nb, KO.VP1-loop-Nb, KO, measured by the percentage of positive GFP cells by flow cytometry, upon infecting HEK293T cells at a constant MOI of 50. Data given as the mean ± s.d. of $n = 3$ replicates. c) Fluorescence images associated with panel b.

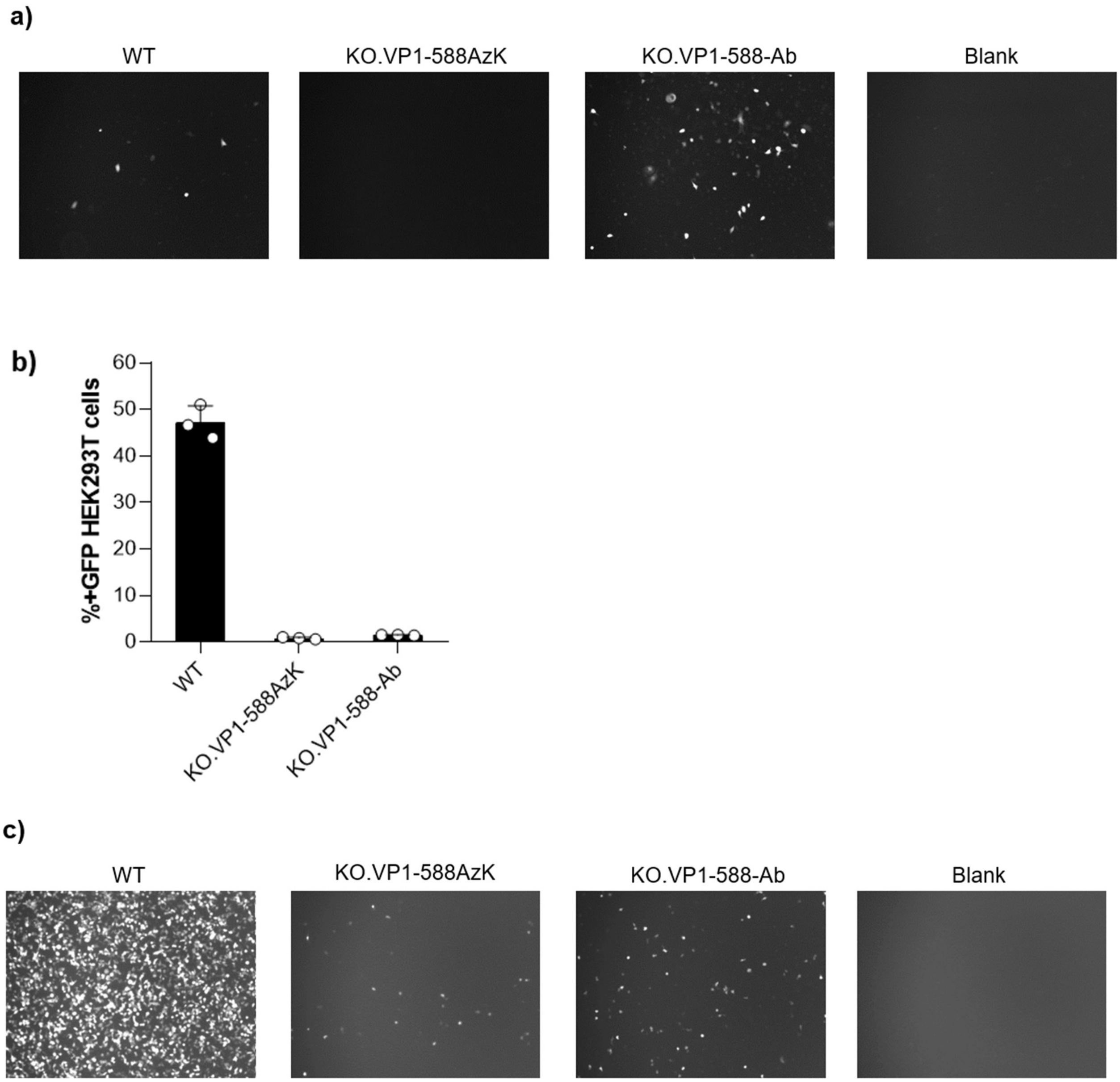


**Figure S12**: a) Representative fluorescence images of cells associated with the experiment described in Figure 4c. SKBR3 cells were infected with a constant MOI 250 of AAV2 WT or AzK mutant virus without (KO.VP1-588-AzK) or with conjugation with AntiHer2-Nanobody (KO.VP1-588-Ab), and imaged 48 h post-infection. b) Infectivity of AAV2-WT, KO.VP1-588-AzK, KO.VP1-588-Ab, measured by the percentage of positive GFP cells by flow cytometry, upon infecting HEK293T cells at a constant MOI 50. Data given as the mean ± s.d. of $n = 3$ replicates. c) Fluorescence images associated with panel b.

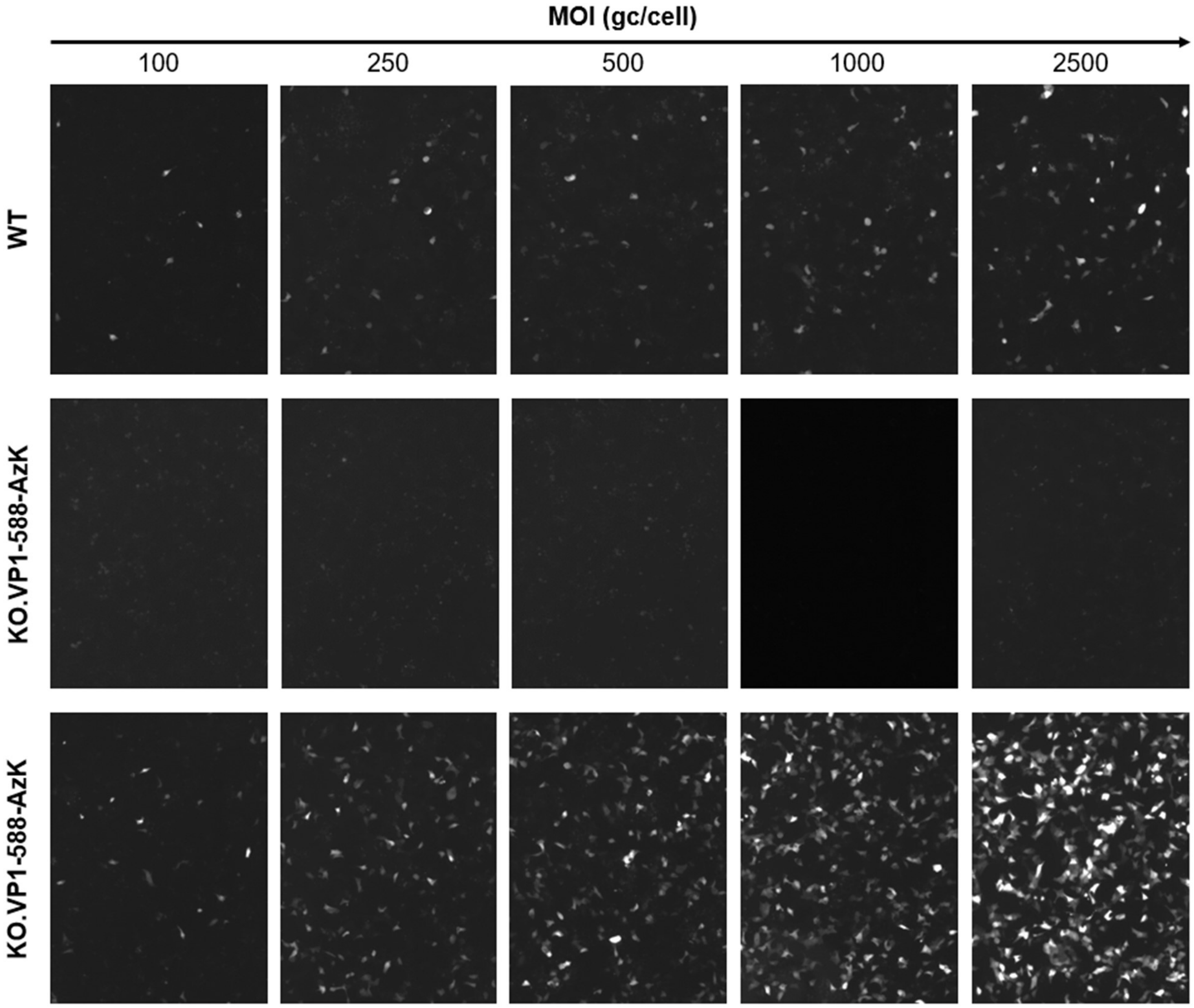


**Figure S13**: Representative fluorescence images of cells associated with the experiment described in Figure 5a. BT474 cells were infected with various MOI (100 – 2500) of AAV2 WT, detargeted virus without (KO.VP1-588-AzK) or with conjugation with AntiHer2-Nanobody (KO.VP1-588Nb), and imaged 48 h post-infection.

a

**Day 14**

Ventral | Dorsal

**Group 1**
WT
(mouse #1-8)

**Group 2**
KO.VP1-588AzK
(mouse #9-16)

**Group 3**
KO.VP1-588AzK
(mouse #17-24)

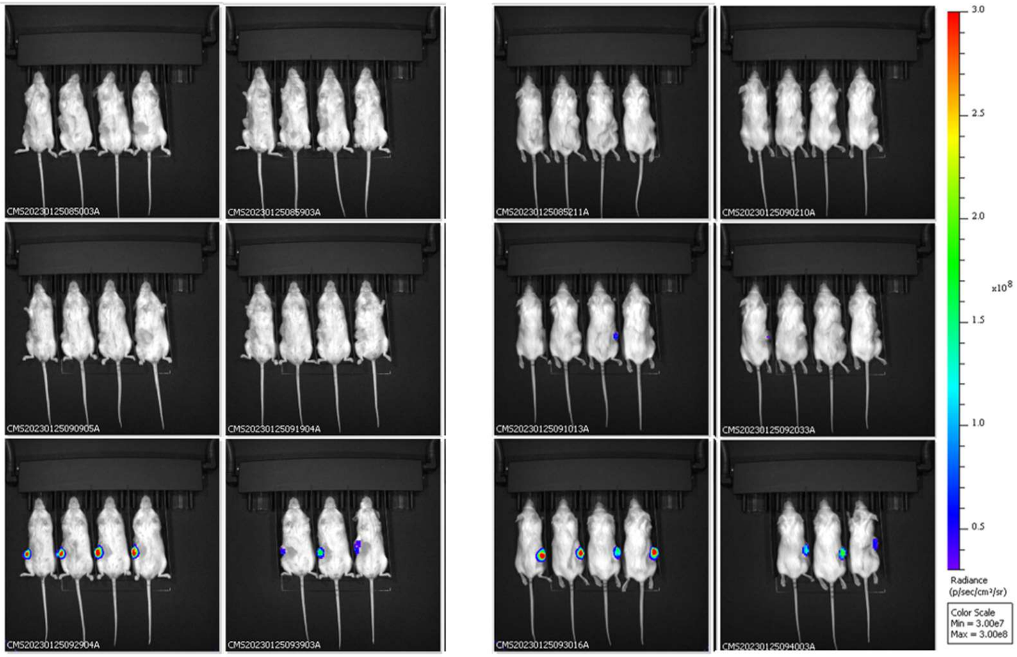


**Day 21**

Ventral | Dorsal

**Group 1**
WT
(mouse #1-8)

**Group 2**
KO.VP1-588AzK
(mouse #9-16)

**Group 3**
KO.VP1-588AzK
(mouse #17-24)

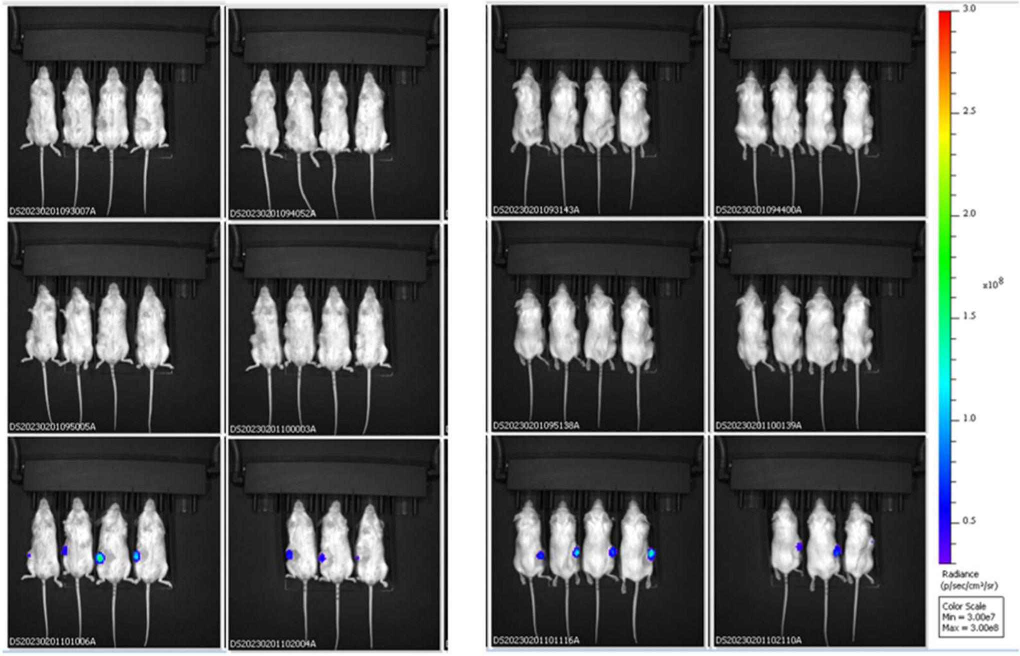


**(continued on next page)**

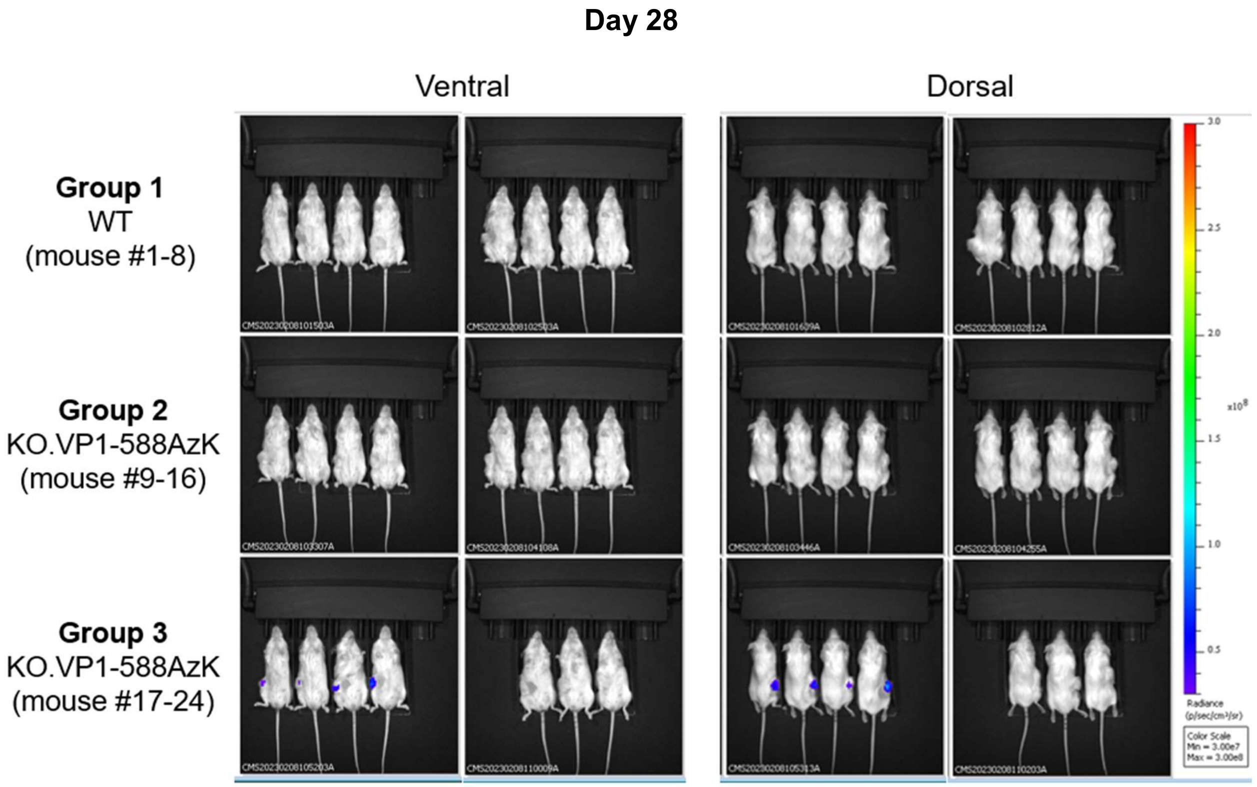


b

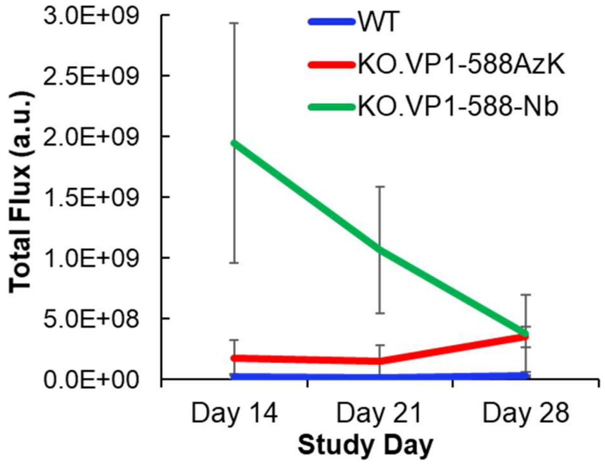


**Figure S14:** a) Ventral and dorsal images of mice treated with AAV2-WT detargeted virus without (KO.VP1-588-AzK) or with conjugation with AntiHer2-Nanobody (KO.VP1-588Nb) throughout the animal study (fixed luminescence scale). Mouse #22 was found dead at day 12 with no obvious cause of death. b) Average total flux obtained from whole-body fluorescence imaging from the three different animal groups over the 5-week study period.

#### Materials and Methods

**Cell Culture.** HEK293T cells, Expi293 were obtained previously[1,2]. SK-BR-3 cells were a gift from Prof. Eranthie Weerapana, Boston College Chemistry, and BT-474 cells was a gift from Charles River Laboratory. All cell lines were cultured at 37 °C and 5% CO2 in DMEM-high glucose (HyClone) supplemented with penicillin/streptomycin (HyClone, final concentration of 100 U/mL penicillin and 100 μg/mL streptomycin) and 10% fetal bovine serum (Corning). Expi293 cells were cultured at 37 °C, 8% CO2, in Expi293™ Expression Medium (Gibco) supplemented with 0.5x antibiotic-antimycotic (Thermo Fisher) in shaken ventilated Erlenmeyer flasks at 125 rpm.

**Cloning and Plasmids.**

Transformation was done in Top10 cells using BioRad electroporator. DNA oligo synthesis and Sanger Sequencing were performed by Genewiz. Phusion polymerase was purchased from Thermo Scientific™, PrimeSTAR® Max DNA Polymerase was purchased from Takara, T4 DNA Ligase was purchased from Qiagen.

pET22b-T5-sfGFP-151TAG was obtained previously[3]. pET22b-T5-5F7-69TAG was generated by QuickChange site-directed mutagenesis from pET22b-T5-5F7-69TGA[2] using 5F7-69TAG-F and 5F7-69TAG-R primers. pcDNA3.1-Her2-Ab-LC-122TAG was generated by overlap extension PCR on pcDNA3.1-Her2-Ab[2] using pcDNA-LC-SpeI-F, HerAb-LC-122TAG-R, HerAb-LC-122TAG-F, pcDNA-LC-SbfI-R primers. The PCR product was digested and cloned into pcDNA3.1-Her2-Ab using SpeI and SbfI restriction sites.

pIDTSmart-MbPylRS-4xPyOtR-ITR-GFP was generated from original plasmid pIDTsmart-MbPylRS[1]. 4xPyOtR cassette was amplified by PrimeStar polymerase on pIDTsmart-4xPyOtR[4] using NheI-HTS25-F and SbfI-HTS25-R primers. The PCR product was digested with restriction enzymes NheI and SbfI and cloned into pIDTsmart-MbPylRS using SpeI and SbfI restriction sites, yielding the intermediate plasmid pIDTSmart-MbPylRS-4xPyOtR. Next, ITR-EFGP was digested from the plasmid pIDTSmart-8xPyltR-ITR-GFP[1] with restriction enzymes SbfI, and cloned into pIDTSmart-*Mb*PylRS-4xPyOtR using SbfI restriction site. ITR-FLUC was amplified by PrimeStar polymerase from pAAV.CMV.ffLuciferase.SV40 (Addgene) using FLUC-cargo-SbfI-F and FLUC-cargo-SbfI-R primers. The PCR product was digested and cloned into pIDTSmart-MbPylRS-4xPyOtR-ITR-GFP using SbfI restriction site, yielding pIDTSmart-MbPylRS-4xPyOtR-ITR-FLUC.

pHelper, pIDTSmart-RC2-WT, pIDTSmart-RC2-ΔVP1-CMV-VP1, pIDTSmart-RC2-ΔVP12-CMV-VP12 clonings were obtained previously[1]. To detarget AAV from heparan sulfate, two mutations R585A and R588A were made by QuickChange site-directed mutagenesis from pIDTSmart-RC2-WT using 585A/588A-F, 585A/588A-R primers, yielding pIDTsmart-RC2.KO. The resulting plasmid was used as a template to clone pIDTSmart-RC2-ΔVP1-CMV-VP1, pIDTSmart-RC2-ΔVP12-CMV-VP12 with R585A and R588A mutations at all VP1,2,3 as previously described[5]. For the incorporation of ncAA into different sites of VP1, overlap-extension PCR on pIDTSmart-RC2-ΔVP1-CMV-VP1 was done to introduce TAG codon to CMV-VP1 gene using SbfI-CMV-F, Cap2-Bsu36I-R, and suitable overlap primers. The PCR products were digested and cloned into pIDTSmart-RC2-ΔVP1-CMV-VP1 using NheI, Bsu36I restriction sites. For the incorporation of ncAA into different sites of VP1+VP2, overlap-extension PCR on pIDTSmart-RC2-ΔVP2-CMV-

VP2 was done to introduce TAG codon to CMV-VP2 gene using SbfI-CMV-F, Cap2-MluI-R, and suitable overlap primers.

pIDTSmart-RC2-ΔVP1-CMV-VP1-loop-Nb was generated by 2 steps overlap-extension PCR, and cloned into pIDTSmart-RC2.KO-ΔVP1-CMV-VP1 using SbfI and Bsu36I restriction sites. The first overlap-extension PCR was done to fuse CMV-N terminus-VP1 (codons 1-452) with anti-Her2 Nanobody (clone 5F7) gene, flanked by 5xGGGGS flexible linker using CMV-SbfI-F, VP1half-GS-R, GS-5F7-F, 5F7-GA-Bsu-R primers. The second overlap-extension PCR was done to fuse CMV-N terminus-VP1-Nb with C terminus-VP1 genes (codons 450-735) using CMV-SbfI-F, VP1-2$^{nd}$ half-F, 5F7-overlapVP-R, Cap2-Bsu36I-R primers. pIDTSmart-RC2-ΔVP2-CMV-5F7-VP2 was generated by overlap-extension PCR to fuse anti-Her2 Nanobody to N terminus of VP2, flanked by 1xGGGGS linker, using NheI-5F7-F, 5F7-VP2 overlap-F, 5F7-VP2 overlap-R, Cap2-Bsu36I-R primers. The PCR product was digested and cloned into pIDTSmart-RC2-ΔVP2-CMV-VP2 using NheI and Bsu36I restriction sites.

**Primers used in this study**:

| Primer name | Sequence |
|---|---|
| 5F7-69TAG-F | ggccgcttttagattagccgcgataacgcgaaaaacac |
| 5F7-69TAG-R | cgcggctaatctaaaagcggcctttcacgctatc |
| pcDNA-LC-SpeI-F | cctggagacgccatcacactagtcaccatgagggtccccgc |
| pcDNA-LC-SbfI-R | agtgacctgcaggttattaacactctcccctgttgaagctctttg |
| HerAb-LC-122TAG-F | cttcatcttcccgccatcttaggagcagttgaaatctggaac |
| HerAb-LC-122TAG-R | caactgctcctaagatggcgggaagatgaagacagatggtgc |
| NheI-HTS25-F | gtttgagacgggcgacagatc |
| SbfI-HTS25-R | atattaattcctgcaggctcgatccgctcgcaccc |
| FLUC-cargo-SbfI-F | aacccttggaatacctgcaggattaaggccttaattagg |
| FLUC-cargo-SbfI-R | aattaaaattcctgcaggcgacggccagtgaattag |
| 585A/588A-F | cctccaggccggcaacgcccaagcagctaccgcagatgtcaacacac |
| 585A/588A-R | gcttgggcgttgccggcctggaggttggtagatacagaaccatactgctc |
| SbfI-CMV-F | aggtaccttcaacctgcaggttgacattgattattgactag |
| Cap2-Bsu36I-R | tcagtattgagcctcaggtgtagttaatgattaacccgccatgctacttatc |
| Cap2-MluI-R | taagagaattacgcgttgtagttaatgattaacccgccatgc |
| Q263TAG-F | caaacaaatttccagctagtcaggagcctcgaacgacaatcac |
| Q263TAG-R | cgaggctcctgactagctggaaatttgtttgtagaggtggttgttg |
| T456TAG-F | gtggaaccacctagcagtcaaggcttcagttttctcaggccg |
| T456TAG-R | gaagccttgactgctaggtggttccacttggagtgtttgttc |
| R588TAG-F | ctgcggtagctgcttgctagttgcctctctggaggttggtagatacag |
| R588TAG-R | ccagagaggcaactagcaagcagctaccgcagatgtcaacacac |
| VP1half-GS-R | ccacctgatccaccacctcctgaacctcctcctcctgaacctcctcctccacttggagtg |
| GS-5F7-F | ggaggtggtggatcaggtggcggtggttctggaggaggaggttctgaagtgcagctggtg |
| 5F7-GA-Bsu-R | attgagcctcaggggctccaccgccaccgctgctcacggtcac |
| NheI-5F7-F | aagctggctagcatggaagtgcagctggtgg |

| | |
|---|---|
| 5F7-VP2 Overlap-R | cggagcggtggatccaccgccaccgctgctcacggtcacttg |
| 5F7-VP2 overlap-F | gcagcggtggcggtggatccaccgctccgggaaaaaagaggc |

**Production of ncAA-containing AAV**

AAV2 was produced by transfecting HEK293T cells with pHelper, pIDTsmart-*Mb*PylRS-4xPyOtR-ITR-GFP, and plasmid containing suitable AAV Rep-Cap genes in 1:1:1 molar ratio[5]. The DNAs were premixed in serum free media (DMEM) and PEI (sigma) was added (3.5 µL per µg DNA). The mixture was allowed to sit at RT for 15 minutes before adding dropwise to the cells, followed by the addition of AzK (Iris Biotech) to the final concentration of 1 mM. 57 µg total DNAs were used for a 15 cm dish and viruses were harvested 120 hours post transfection. Virus was extracted from cells by 2 cycles of snap freeze-thaw in dry ice-ethanol and 37C water bath. Cell lysate was combined with media and was mixed with 30% volume of 40% PEG 8000 (Fisher BioReagents) to precipitate the virus overnight at 4 °C. On the next day, virus was spun down at 5,000xg for 30 minutes, then resuspended in 2 mL of salty PBS (300mM NaCl), supplemented with 2 µL universal nuclease and nutated at RT for 15 minutes. During that time, 250 µL slurry of AVB-agarose was packed in a column and equilibrated with 10x column volume of dPBS. Virus was loaded to the column and flowed down by gravity and then reloaded twice. The column was washed with 30x column volume of dPBS. Virus was quickly eluted by 2 mL 0.1 M Glycine pH 2.8 and neutralized with 200 µL 1M Tris pH 8. Virus was concentrated and buffer-exchanged to AAV buffer (1xdPBS, 300 µM NaCl, 0.001% Pluronic F68) using an Amicon 100 kDa molecular weight cut-off centrifugal filter. Glycerol was added to the final concentration of 10% and virus was titered by AAVpro® Titration Kit Ver.2 (Takara), followed the manufacturer's instructions. For animal studies, the viruses were sterile filtered by passing through a 0.2 µm, 4 mm filter. The content of endotoxin in the virus samples were assayed and found to be <0.900 EU/ml using the ToxinSensorTM Chromogenic LAL Endotoxin Assay Kit (Genscript).

**Protein expression**

sfGFP-151-AzK[3], AntiHer2-Nanobody (Clone 5F7)-69-AzK[2], and full length Trastuzumab-LC-122-AzK[6] expression was done using the previously described methods.

For animal study, AntiHer2-Nanobody was sterile filtered by passing through a 0.2 µm, 4 mm filter. Endotoxin was removed from AntiHer2-Nanobody sample by Pierce™ High Capacity Endotoxin Removal Spin Columns, Capacity = 0.5 mL (Thermo Scientific), followed the manufacturer's instructions.

**Protein-protein conjugation**

GFP coupling: 100 µM sfGFP-151AzK and 50 mM DBCO-PEG4-DBCO (Broadpharm) or DBCO-sulfo-Tz (Broadpharm) or DBCO-PEG4-TCO (Broadpharm) were diluted in 1% DMSO in DPBS (pH = 7.4) to a final concentration of 10 µM and 100 µM, respectively. Conjugations were done in 50 µL scale in 0.5 mL tubes by nutating the mixtures at 22°C for 12h. The modified proteins were buffer exchanged in DPBS using Amicon 10 kDa molecular weight cut-off centrifugal filters device to remove excess DBCO probes. sfGFP containing DBCO and sfGFP-151-AzK concentration were adjusted to 10 µM and mixed together in 1:1 molar ratio. GFP coupling was done by nutating the mixture at RT for 16 hours. sfGFP containing TCO and Tz concentration were adjusted to 10 μM and mixed together in 1:1 molar ratio. GFP coupling was done by nutating the mixture at RT and quenched by the addition of 50-fold excess DBCO-sulfo-Tz.

**Nanobody-virus conjugation**: 100 μM AntiHer2-Nb-69-AzK and 50 mM DBCO-PEG4-TCO or DBCO-PEG12-TCO (Broadpharm) were diluted in 1% DMSO in DPBS (pH = 7.4) to final concentration of 10 μM and 100 μM, respectively. Conjugations were done in 200 μL scale in 0.5 mL tubes by nutating the mixtures at 22°C for 16h. The modified proteins were buffer exchanged in DPBS using ultracentrifugal device 0.5 mL 10kDa to remove excess DBCO probes. 50 mM DBCO-sulfo-Tz or DBCO-PEG12-Tz (Broadpharm) were diluted in AzK-containing virus solutions to a final concentration of 100 μM. Reactions were done in 1.5 mL tubes by nutating the mixtures at 22°C for 16h, before complete buffer exchange into AAV buffer by ultracentrifugal device Amicon 100 kDa to remove excess DBCO probes. Virus containing Tz was retitered and nanobody containing TCO was added to the final concentration of 1 μM. Conjugation was done in 1.5 mL tubes by nutating the mixtures at 22°C for 4 h.

**Full-length Antibody-virus conjugation**

5 mg/mL Trastuzumab-LC-122-AzK and 50mM DBCO-PEG12-TCO were diluted in 1% DMSO in DPBS (pH = 7.4) to final concentration of 1 mg/mL and 100 μM, respectively. Conjugations were done in 200 μL scale in 0.5 mL tubes by nutating the mixtures at 22°C for 16h. The modified proteins were buffer exchanged in DPBS using ultracentrifugal device 0.5 mL 50 kDa to remove excess DBCO probes. 50 mM DBCO-PEG12-Tz was diluted in AzK-containing virus solutions to a final concentration of 100 μM. Reactions were done in 0.5 mL tubes by nutating the mixtures at 22°C for 16 h, before complete buffer exchange into AAV buffer by ultracentrifugal device Amicon 100 kDa to remove excess DBCO probes. Virus containing Tz was retitered and Antibody containing TCO was added to the final concentration of 0.5 μM. Conjugation was done in 0.5 mL tubes by nutating the mixtures at 22°C for 4h.

**LC-MS analysis**

LC-MS was done using Agilent Technologies, 1260 Infinity ESI–TOF, Phenomenex, Aeris™ 3.6 μm WIDEPORE XB-C8 column, LC Column 100 x 4.6 mm. Separation was performed with a flow rate of 0.2 mL/min, and with mobile phase of solvent A (water-MeCN-0.1% trifluoroacetic acid in 95:5:0.1 ratio) and solvent B (MeCN-water-0.1% trifluoroacetic acid in 95: 5: 0.1 ratio); 0-1 min 5% B, 1-8 min 99% B, 8-9 min 99% B to 1% B, 9-10 min 1% B to 5% B, 10-14 min 5% B. Deconvolution was done using Magtran software, and peak intensities for Azido-lysine containing proteins and their corresponding linker conjugated products were compared.

For the LC-MS analysis of Trastuzumab-LC-122-AzK and DBCO-PEG12-TCO adduct, the antibodies were reduced with 5 mM DTT at 55°C for 10 minutes. The reduced antibodies (25 μL, 1 mg/mL) were treated with 1 μL Remove-iT® PNGase F (NEB) for 2 hours at 37°C to remove the antibody glycosylation. Due to the instability of the TCO during Antibody treatment for mass spectrometry, Trastuzumab-LC-122-AzK and DBCO-PEG12-TCO adduct was reacted 200 μM Tetrazine-aniline for 10 minutes at RT prior to LCMS analysis.

**SDS Page**

Approximately 5 µg of sfGFP and $10^{10}$ genome copies of viruses were loaded per lane. The samples were heated in SDS-loading buffer for 1 min, then analyzed with 10% SDS-PAGE gel. sfGFP were stained with Commasie brilliant Blue and imaged on the ChemiDoc MP (BioRad) under Commassie blue setting. Viruses and attached proteins were stained with SYPRO™ Orange Protein Stain (Thermo Fisher Scientific), followed the manufacturer's instructions, and imaged with Dylight 540.

**Assaying the infectivity of AAV2 *in vitro* and flow cytometry.**

Infectivity in HEK293T cells was assayed as previously described[1]. For SKBR3 cells, 2.5 x$10^6$ cells (counted by Bio-Rad TC20™ Automated Cell Counter) were seeded per 24-well plate 30 hours prior to infection. For BT474 cells, 5.0 x$10^6$ cells were seeded per 24-well plate 24 hours prior to infection. When cells had reached the desired confluency, viruses were added to each well in a constant MOI, along with 5 mM sodium butyrate (Sigma-Aldrich) to enhance the expression of AAV2-encoded transgenes. 48 h post-infection, media was removed and 200 µL RT DPBS was added per well. Infectivity was visualized by EGFP expression using a Zeiss Axio Observer fluorescence microscope with an XCite Series 120Q light source and Zeiss filter 44 (excitation 475/40 nm, beamsplitter 500 nm, emission 530/50 nm). Cells were harvested by 5 incubation in 100 µL warm 0.25% trypsin-EDTA at 37$^o$C, followed by quenching with 200 µL of cold DMEM supplemented with 10% FBS. Cells were resuspended and pelleted in 2 mL eppi tubes by centrifugation at 2,500 xg for five minutes at 4$^o$C. The supernatant was discarded and cells were gently resuspended in 200 µL of cold PBS and passed through a 35 µm strainer cap of a 5 mL tube (Falcon). Flow cytometry was performed on a BD Accuri C6 Plus flow cytometer (BD Biosciences) using the FITC and PerCP settings (excitation wavelength: 488 nm, standard filter: 533/30, 670LP, respectively). Data were processed using FlowJo, version 10.8.1.

#### Representative examples of gating and FACS analysis

Gating example of SK-BR-3 cells

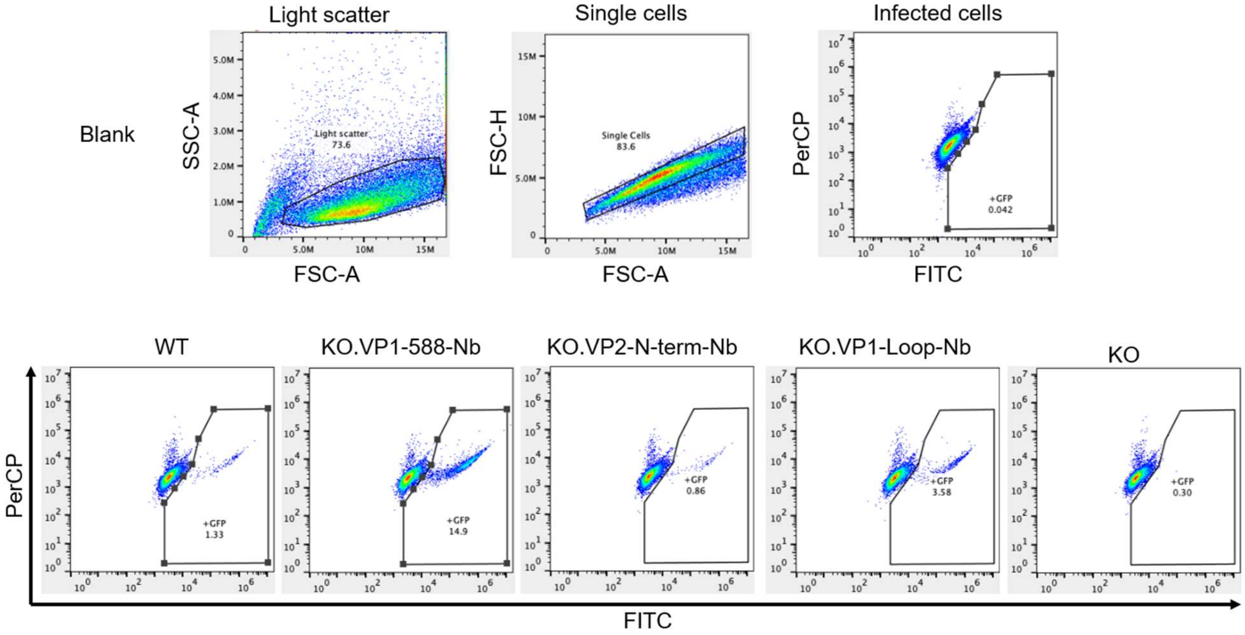


Gating example of HEK293T cells

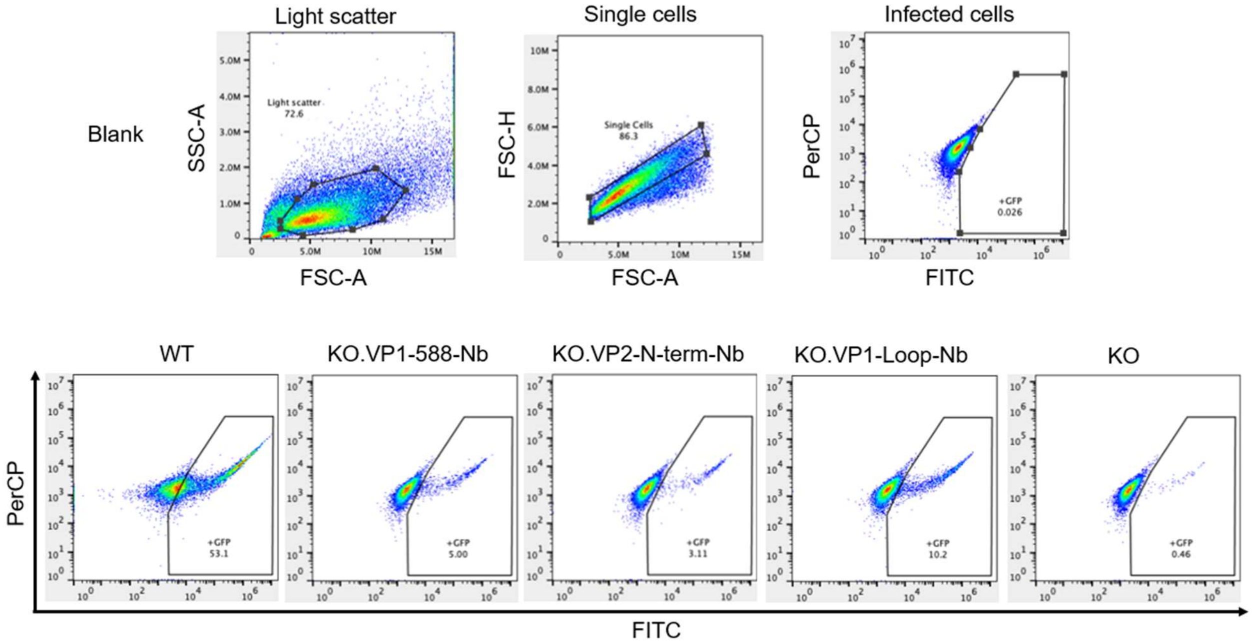

**Animal studies, in vivo imaging, and transgene titering.**

Animals were prepared and *in vivo* imaging was performed by Charles River Laboratories (Worcester, MA). 40 female mice (8 weeks old) were xenografted with 5.0 x$10^6$ BT-474 cells. Tumors were allowed to expand and 8 mice were randomly pooled such that the average tumor per group volume is 100 $mm^3$. 1.0 x $10^{11}$ gc each of WT, KO.VP1-588-AzK, and KO. VP1-588-Nb, encoding the firefly luciferase reporter was used to infect the animals by tail-vein injection. Starting on day 14 post-injection, luminescence was measured weekly using the IVIS Lumina III imaging system. Animals were sacrificed at day 35 and tumors, livers were collected and stored in RNA later. The tissue samples were ground and homogenized in a BeadbugTM3 microtube homogenizer, 115V (Benchmark Scientific) using prefilled 2.0 mL tubes with impact zirconium Beads (Benchmark Scientific). Total DNA was isolated from about 25 mg tissue sample using DNeasy Blood & Tissue Kit (Qiagen), following the manufacturer's instructions. DNAs were ethanol precipitated for an hour and resuspended in milliQ water. AAV transgene was titered by AAVpro® Titration Kit Ver.2 (Takara), following the manufacturer's instructions.

(4) Jewel, D.; Kelemen, R. E.; Huang, R. L.; Zhu, Z.; Sundaresh, B.; Malley, K.; Pham, Q.; Loynd, C.; Huang, Z.; van Opijnen, T.; Chatterjee, A. Enhanced Directed Evolution in Mammalian Cells Yields a Hyperefficient Pyrrolysyl tRNA for Noncanonical Amino Acid Mutagenesis. *Angewandte Chemie International Edition n/a* (n/a), e202316428. https://doi.org/10.1002/anie.202316428.

(5) Erickson, S. B.; Pham, Q.; Cao, X.; Glicksman, J.; Kelemen, R. E.; Shahraeini, S. S.; Bodkin, S.; Kiyam, Z.; Chatterjee, A. Precise Manipulation of the Site and Stoichiometry of Capsid Modification Enables Optimization of Functional Adeno-Associated Virus Conjugates. *Bioconjugate Chem.* **2024**, *35* (1), 64–71. https://doi.org/10.1021/acs.bioconjchem.3c00411.

(6) Zhao, C.; Wang, Y.; Pham, Q.; Dai, C.; Chatterjee, A.; Wasa, M. Chemical Tagging of Bioactive Amides by Cooperative Catalysis: Applications in the Syntheses of Drug Conjugates. *J. Am. Chem. Soc.* **2023**, *145* (26), 14233–14250. https://doi.org/10.1021/jacs.3c00169.
